# The transcriptional response of *Yersinia pseudotuberculosis* to macrophage-released metabolites during growth within synthetic microcolonies

**DOI:** 10.64898/2026.03.25.714363

**Authors:** Stacie A. Clark, Alexander Palmer, Wenwen Huo, Alexander C. Joyce, Kimberly M. Davis, Juan Ortiz-Marquez, Tim van Opijnen, Ralph R. Isberg

## Abstract

*Yersinia pseudotuberculosis (Yptb)* replicates in immune cell-encompassed microcolonies within tissues. Bacterial replication is controlled by protection against neutrophil attack and by macrophage-released antimicrobial factors, such as nitric oxide (NO). During these attacks, bacteria located on the microcolony periphery encounter extracellular signals that differ from those in the interior. To dissect individual microbial populations, γ interferon-activated macrophages were used to challenge microdroplet-grown *Yptb* harboring an NO-responsive mCherry reporter. Subsequently, bacterial subpopulations that hyperactivated the reporter were isolated from droplets composed of a reversible polymer matrix. RNA-seq analysis indicated that induction of nitrosative stress-associated genes was the primary determinant distinguishing peripheral bacteria from the remaining population. In addition, a secondary stress response that induced prophage-associated genes was detected, which could not be traced to either DNA damage or nitrosative stress responses. Activated macrophages also induced the expression of the *Yptb* itaconate degradation enzyme-encoding transcript throughout the entire colony. To determine if itaconate production by the interferon-activated Irg1 protein played a role in restricting *Yptb*, bacteria harboring an itaconate-responsive reporter and *Yptb* mutants defective for itaconate degradation were analyzed during bacterial colonization of the murine spleen. Only a subset of colonies appeared to be exposed to itaconate, which may explain the very small defects exhibited by mutants unable to degrade the interferon-induced macrophage product. These results indicate that the primary response of bacteria to macrophage-elicited factors is likely associated with protection against NO-derived metabolites.

## Introduction

During systemic infection, bacterial pathogens can colonize and replicate in deep tissue sites, causing acute or chronic disease (1–3). The colonization site is often surrounded by innate immune cells, which can develop into either abscesses, granulomas, or microcolonies, the latter of which have been referred to as pyogranulomas (4–7). *Yersinia pseudotuberculosis* (*Yptb*) is one such pathogen that establishes disease by forming microcolonies within deep tissues sites during disease (3, 8). After intestinal inoculation of mice, microorganisms replicate in the intestinal lumen and regional lymph nodes, and disseminate via a poorly characterized route into the liver or spleen (9).

During microcolony formation in these deep tissue sites, *Yptb* establishes clonal extracellular microcolonies that develop into lesions densely populated with innate immune cells (3, 8) (10). Neutrophils are likely the first immune cells recruited to the focus of replicating bacteria, directly surrounding the growing microcolony. *Yptb* in the periphery of the microcolony are in direct contact with neutrophils, resulting in the translocation of the bacterial Type III Secretion System (T3SS) effector proteins into target host cells, paralyzing cytoskeletal elements and misregulating components of the innate immune response (11–16). As a result, the pathogen disrupts phagocytosis and the production of reactive oxygen species (ROS) by surrounding neutrophils (17). The paralyzed neutrophils, therefore, are unable to limit Yptb growth, likely leading to the subsequent recruitment of additional innate immune cells, such as macrophages and monocytes. This second wave of recruited immune cells surrounds the sphere of neutrophils, resulting in a complex, multilayered structure.

The recruitment of myeloid cells around a cluster of *Yptb* exposes the bacterial microcolony to distinct microenvironments, which drives the phenotypic specialization of microbial subpopulations that counter host attack. Peripheral bacteria in direct contact with neutrophils upregulate their T3SS, amplifying the anti-host cell response and rendering the ring of neutrophils ineffective in clearing the bacteria (8). The cell-associated *Yptb* also contributes to a larger peripheral subpopulation of bacteria that upregulate the protein Hmp that detoxifies nitric oxide (NO) gas produced by surrounding macrophages. Consequently, the majority of the peripheral cells responding to NO do not come into direct contact with any cells, and few, if any, of the NO-producing host cells are in direct contact with the microcolony. The bacteria at the center of the microcolony are protected from NO by the Hmp-expressing peripheral bacteria, providing an example of tissue-associated microbial social behavior in which one subpopulation protects the inner core. The ability of *Yptb* to detoxify NO *via* Hmp is important for virulence, as microcolonies lacking Hmp eventually disintegrate in tissues (8).

Until recently, analysis of bacterial subpopulations was limited to microscopic analysis of fixed tissue or tissue culture modeling strategies that rely on microbial challenge of cells incubated in 2-dimensional culture. Analysis of fixed tissue provides a limited opportunity to perform genetic analysis, and incubations with mammalian cultured monolayers do not reproduce the three-dimensional (3D) architecture of bacterial microcolonies surrounded by immune cells in host tissue sites. To bridge the gap between animal infection models and tissue culture models, we previously developed an *in vitro* system that uses droplet-based microfluidics to generate matrix-embedded *Yptb* microcolonies, incorporating primary macrophages that bind to the outside of the droplet (18). Social behavior in response to macrophages could be reproduced by an NO generator or by macrophages activated to produce inducible nitric oxide synthase (iNOS). The significance of this model is that topological analysis of *Yptb-*host cell interaction is unattainable in murine and tissue culture models of infection. It remains unclear what the growth dynamics of bacterial subpopulations are in this micro-compartment, or if other macrophage-secreted factors are being produced by macrophages that target *Yptb*.

In this report, we modified the microdroplet culture strategy to efficiently isolate bacterial subpopulations *via* flow cytometry and perform transcriptional analysis on the *Yptb* subpopulations. Similar to our original system, social behavior was reproduced in the presence of a chemical NO generator or by activated macrophages expressing iNOS. Furthermore, we also provide evidence that the macrophage metabolite itaconate, synthesized as part of the interferon response, is released into the bacterial microenvironment, and may contribute to control of *Yptb* growing extracellularly in tissue sites.

## Results

### Alginate microdroplets support growth of *Yersinia pseudotuberculosis* microcolonies

*Yptb* forms clonal microcolonies in the murine spleen that replicate extracellularly and are surrounded by innate immune cells (3, 8). This 3-dimensional (3D) topology can be mimicked in culture in which a matrix consisting of 1% ultra-low melt agarose containing 25% HyStem^®^-C Hydrogel encapsulates *Yptb* in 65 μm-diameter droplets (18). This matrix requires heating to isolate bacteria for sorting populations based on fluorescent protein expression. Unfortunately, the chemicals used to inactivate RNase activity resulted in a loss of protein fluorescence during the heating step (Supplemental Fig. 1). To address this problem, bioengineered droplets were prepared using alginate, which enables EDTA dissolution of the droplets, thereby avoiding a heating step (19–22). Alginate is a polysaccharide consisting of guluronate and mannuronate chains (23). The carboxylate groups of these residues bind most divalent cations, most commonly Ca^2+^, thereby allowing the alginate to form a gel immediately.

A strategy was devised to enable controlled gelation of alginate droplets using microfluidics, thereby avoiding device clogging (24). To allow controlled gelation, Ca^2+^/EDTA chelation followed by EDTA acidification is often employed (25), but this strategy was not effective because EDTA blocked bacterial growth in droplets, as reported previously (26, 27). As an alternative, dispersed CaCO_3_ nanoparticles were incorporated into the system, followed by acidification after microdroplet formation to solubilize Ca^2+^. To this end, 25 mM CaCO_3_ nanoparticles were added to a matrix consisting of molten 1% alginate, with the integrin-binding RGD peptide linked to alginate as 5% of the polymer content, to allow multivalent adhesion to mammalian integrin receptors (Fig. 1) (18, 19, 28–32). The mixture was then encapsulated into droplets in oil using a microfluidics chip (18), and the alginate was crosslinked by adding dilute acetic acid in oil (Fig. 1A: step 1-3). After crosslinking (Fig. 1A: step 4), the oil was removed, yielding alginate droplets of approximately 65 μm in diameter (∼144 pl) (Fig. 1B).

**Figure 1.**
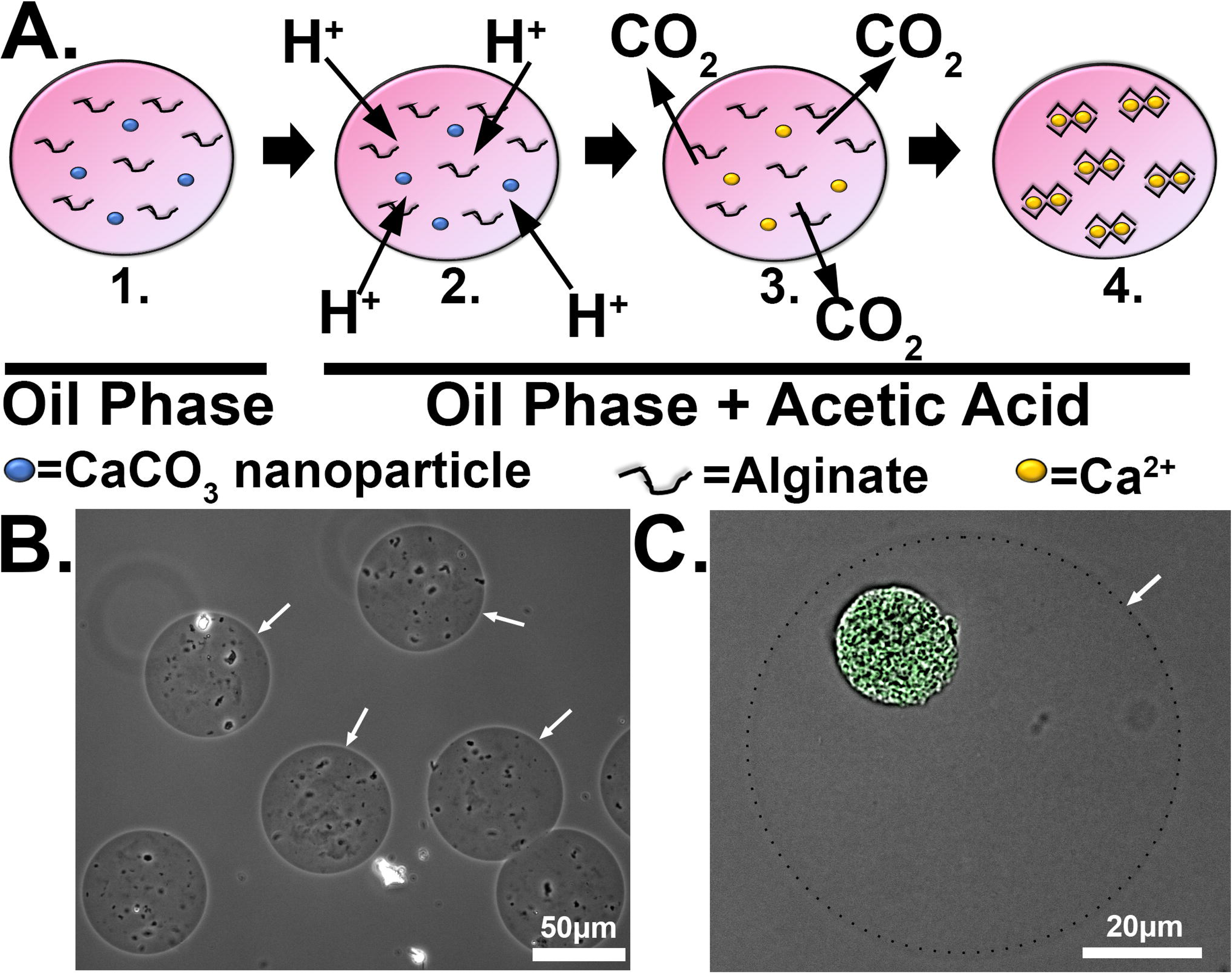
*Yersinia pseudotuberculosis* grows as a microcolony in microfluidics-driven alginate droplets. (A). Construction of alginate droplets. 1) Molten alginate, of which 5% is RGD-linked, and 25 mM CaCO_3_ nanoparticles were introduced into a microfluidics device, allowing encapsulation within oil-coated droplets. 2) Acetic acid (final concentration 0.05%) was added to alginate-CaCO_3_ droplets in oil to allow for gelation of alginate. Protons diffuse into the alginate-CaCO_3_ droplets in oil. 3) H^+^ ions exchange with Ca^2+^ ions, releasing CO_2_. 4) Free Ca^2+^ binds alginate and crosslinks the matrix. (B). Image of alginate droplets with oil removed after in-oil polymerization of alginate. Scale bar: 50μm. (C). *Y. pseudotuberculosis* grows as a microcolony in droplets. Droplets containing encapsulated *Y. pseudotuberculosis gfp+* were cultured overnight at 26°C, and microcolonies were visualized by brightfield and fluorescence microscopy. Dashes: define outline of droplet. Arrows: edge of droplets. Scale bar: 20μm.

*Yptb* microcolonies form as clusters in tissue (3, 18). To confirm *Yptb* grows as a cluster inside alginate droplets, the *Yptb* strain IP2666 expressing GFP (*P_tet_::gfp;* Table 2) was encapsulated in droplets, oil was removed, and the droplets were aerated at 26 °C in broth supplemented with 5 mM CaCl_2_ (Materials and Methods). Microcolonies were established that were indistinguishable from the previously described agarose microcolonies (Fig. 1C) (18). Therefore, a tunable and reversible matrix allows microcolony formation of *Yptb*, mimicking topological constraints observed in tissue.

### Spatial regulation of *hmp* expression in tissues can be reproduced in alginate microdroplets

To determine if the spatial regulation of *Phmp* observed in tissue can be reproduced in alginate droplets, the WT strain carrying the *gfp^+^* and *Phmp::mcherry* reporters was encapsulated in droplets, grown into microcolonies, and exposed to the NO generator DETA-NONOate for 4 hours at 37°C in DMEM (Fig. 2A; Materials and Methods). WT microcolonies showed peripheral expression of *hmp*, while the constitutive GFP signal was uniform throughout the microcolonies, as observed in tissues (Fig. 2B). Using a previously described script to quantitate spatial regulation of *hmp*, the fluorescence at the periphery was calculated relative to the point of lowest intensity (PLI) internal to the microcolony (18). There was a significant increase in the periphery vs. PLI ratio in WT microcolonies in response to DETA-NONOate after 4 hour exposure, indicating peripheral expression of P*hmp::mcherry* (Fig. 2C; P <0.0001), demonstrating that spatial regulation of *hmp* can be reproduced in alginate droplets.

**Figure 2.**
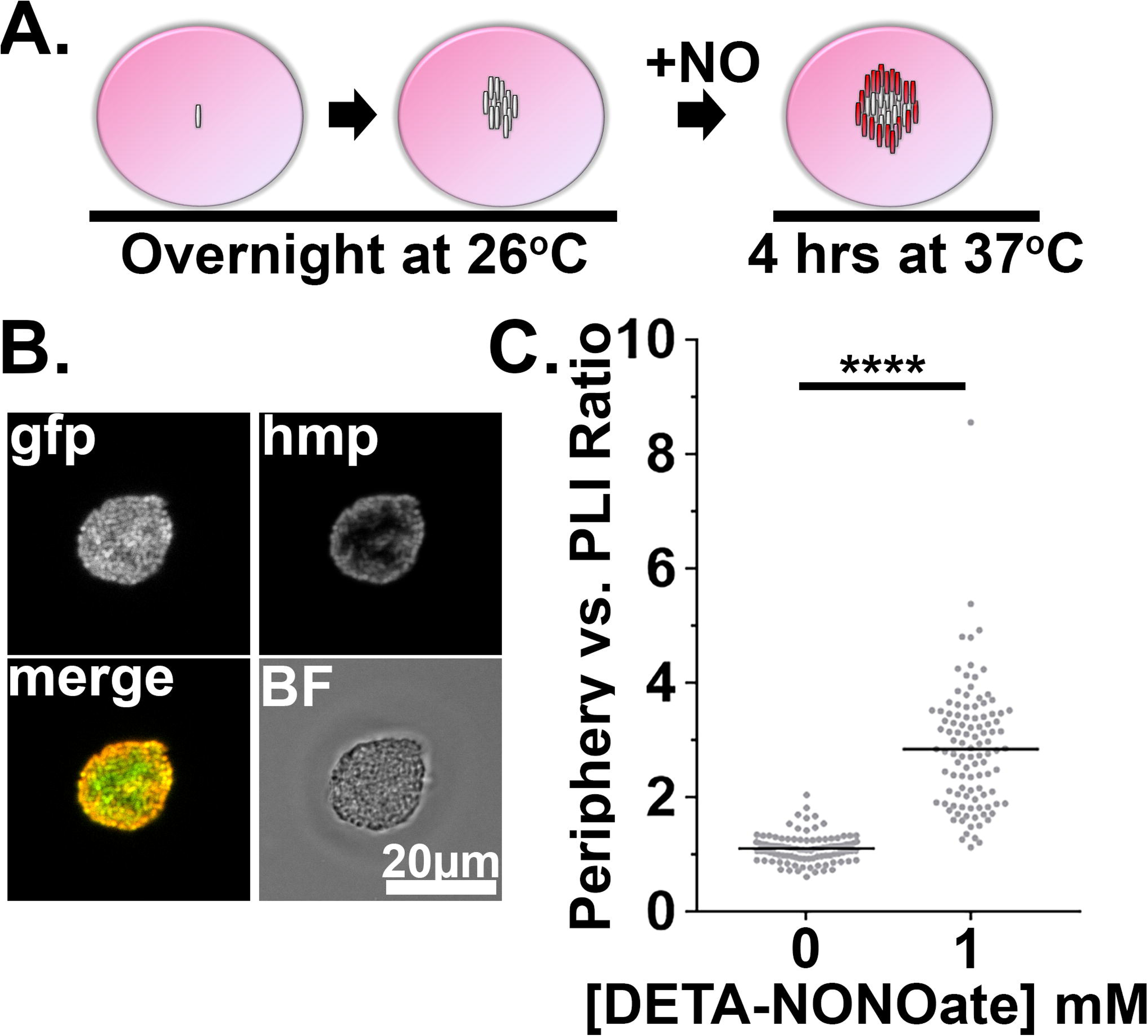
Spatial regulation of *hmp* expression in tissues can be reproduced in alginate droplet culture. (A). Experimental overview. Droplets seeded with *Y. pseudotuberculosis* harboring fluorescence reporters were incubated overnight at 26 °C and 1 mM DETA-NONOate was added to the DMEM cultures for 4 hours at 37 °C, 5% CO_2_. After 4 hours, the droplets were fixed and visualized using brightfield and fluorescence microscopy. (B). Representative WT microcolony after 4 hours of treatment with 1mM DETA-NONOate in DMEM. (C). Periphery versus PLI ratio for microcolonies as a function of DETA-NONOate concentration in microcolonies. Gray circles: 105 individual microcolonies measured for each condition from 3 biological replicates. Lines indicate the medians. Statistics: Mann-Whitney test, ****p<0.0001.

### Activated BMDMs drive peripheral expression of *hmp* in droplet microcolonies

To determine if the microcolonies respond to cellularly produced NO, BMDMs activated by LPS and IFNγ to express inducible NO synthase (iNOS) were added to droplets containing pregrown microcolonies, mimicking tissue dynamics (18, 33–35). BMDMs bound efficiently to alginate droplets containing *Yptb* (Fig. 3A). In response to the primed BMDMs, the *Yptb* microcolonies exhibited spatial regulation of the *Phmp::mcherry* reporter, similar to that observed with *Yptb* growing in tissue (Fig. 3B) (18). Expression of the *Phmp::mcherry* reporter required the addition of LPS and IFNγ to BMDMs, as predicted for driving expression of iNOS (Fig. 3C).

**Figure 3.**
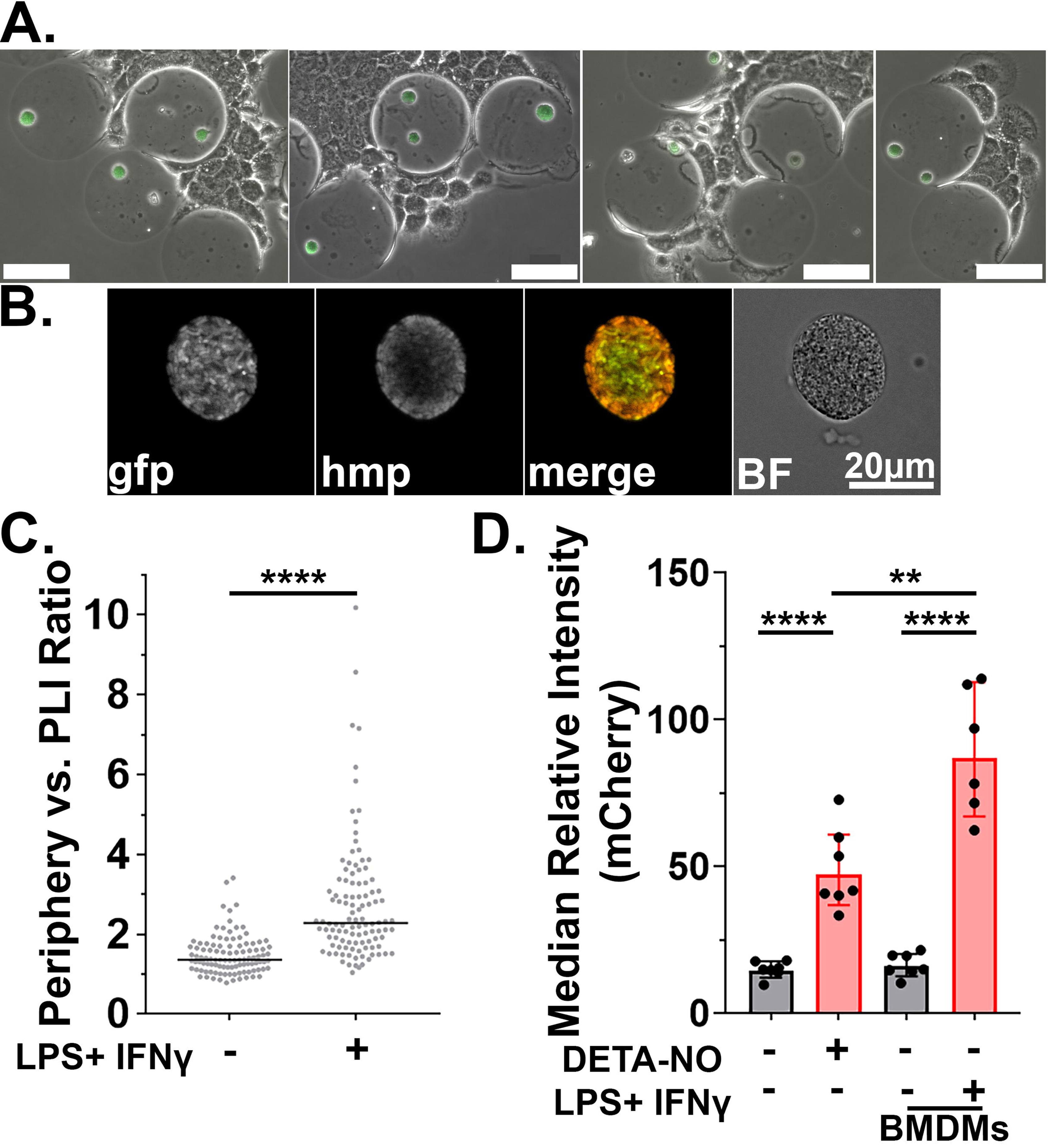
Activated BMDMs drive peripheral expression of *hmp* in alginate droplet microcolonies. (A). BMDMs adhere to alginate droplets containing 5% RGD-alginate. Droplets containing *Y. pseudotuberculosis* were grown overnight in DMEM at 26 °C prior to incubation with LPS/IFNγ-primed BMDMs for 2 hours at 37 °C, CO_2_. The fixed droplets were visualized using phase contrast and fluorescence microscopy. Scale bars: 50μm. (B). Representative WT microcolony incubated with LPS/IFNγ-primed BMDMs in DMEM for 4 hours. Scale bar: 20μm. (C). Periphery versus PLI ratios for microcolonies incubated with BMDMs treated as noted. Gray circles: 105 individual microcolonies from each condition from 3 biological replicates. Lines indicate the median. Statistics: Mann-Whitney test, ****p<0.0001. (D). LPS/IFNγ-primed BMDMs secrete more NO than 1 mM DETA-NONOate. Flow cytometry of dispersed microcolonies after 4 hours of incubation with DETA-NONOate or BMDMs in DMEM. Dispersed bacteria were gated for GFP^+^ and the median mCherry fluorescence intensity was determined from 6-7 biological replicates (Materials and Methods). Lines represent geometric mean with 95% CI. Statistics: Unpaired parametric t test, assuming both populations have the same SD, ****p<0.0001, **p<0.01.

The amount of NO that is generated in tissues surrounding a microcolony is unknown. To determine the relative amount of NO produced by BMDMs, WT microcolonies harboring the *Phmp::mcherry* reporter were challenged in the presence or absence of DETA-NONOate, or unprimed or LPS/IFNγ-primed BMDMs. The droplets were then dispersed by the addition of EDTA and analyzed by flow cytometry, determining the median relative fluorescence intensity of *Phmp::mCherry* per bacterium (Materials and Methods). *Yptb* originating from WT microcolonies incubated with 0 mM DETA-NONOate or unprimed BMDMs, showed low *Phmp::mcherry* fluorescence (Fig. 3D). After challenge with LPS/IFNγ-primed BMDMs, there was a significant increase in *Phmp::mcherry* reporter fluorescence compared to its fluorescence in *Yptb* treated with 1 mM DETA-NONOate (Fig. 3D; p<0.0001). This result is consistent with LPS/IFNγ-primed BMDMs collectively generating more NO than 1 mM DETA-NONOate. Therefore, WT microcolonies exhibit spatial regulation of *hmp* in response to activated BMDMs as seen in tissue, and the relative expression in response to NO secreted by activated BMDMs is more than that observed in response to 1 mM DETA-NONOate.

### Isolation of *Yptb* subpopulations generated in alginate droplets

*Yptb* on the periphery of a microcolony detoxify NO produced by macrophages, lowering NO concentrations below levels necessary to induce *Phmp* in the center of the microcolony. The growth dynamics of these subpopulations within tissue microcolonies is uncertain, and it is not known if there are transcriptionally-distinct bacterial subpopulations controlled by tissue signals. Furthermore, RNA-seq transcriptional analysis of *Yptb* grown in tissue has been limited to whole population analysis without subdividing the response into spatial subpopulations (36). Isolation of bacterial subpopulations from tissue is difficult, and extensive tissue processing may cause significant alterations in the bacterial transcriptome. Therefore, reconstructing spatial regulation in culture is an attractive approach for identifying transcriptional subpopulations. As droplet culture reconstructs the peripheral (NO-detoxifying) and central (non-NO-detoxifying) bacterial subpopulations observed in tissues, it provides a route for analyzing the spatial regulation of these subpopulations in tissue-resident bacteria.

To identify transcriptional differences between *Yptb* peripheral and central populations, *Yptb* subpopulations were fluorescence sorted after growth in alginate droplets. After 7 hours of growth, the WT *gfp^+^/Phmp::mcherry* strain was incubated for an additional 4 hours with either 0 or 1 mM DETA-NONOate, as well as with unprimed or LPS/IFNγ-primed BMDMs. The droplets were then disrupted in the presence of RNA*later™* and sorted (Fig. 4A). Live bacteria were selected by gating for GFP, and peripheral (high Hmp expression; Hmp-HI) versus central populations (low Hmp expression; Hmp-LO) were sorted based on *Phmp::mcherry* fluorescence (Fig. 4B-E). Nearly all WT *Yptb gfp+ bacteria* were Hmp*-*LO without DETA-NONOate (Fig. 4B). After exposure to unprimed BMDMs, most GFP+ bacteria remained Hmp-LO, as expected (Fig. 4D: about 0.5% are Hmp-HI). The slight increase in bacteria within the Hmp*^+^*gate after incubation with unprimed BMDMs compared to 0 mM DETA-NONOate (Fig. 4B, 0.095% Hmp-HI) may result from RGD-alginate activation of the macrophages (37). Conversely, incubation with either 1 mM DETA-NONOate or LPS/IFNγ-primed BMDMs caused strong activation of the *Phmp* promoter in a significant portion of bacteria. To focus on the most highly NO-activated transcripts for transcriptional analysis, GFP+ events were collected from the top 10% of *Phmp::mcherry* fluorescence after treatment with either 1 mM DETA-NONOate (Fig. 4C) or incubation with macrophages treated with LPS/IFN (Fig. 4E).

**Figure 4.**
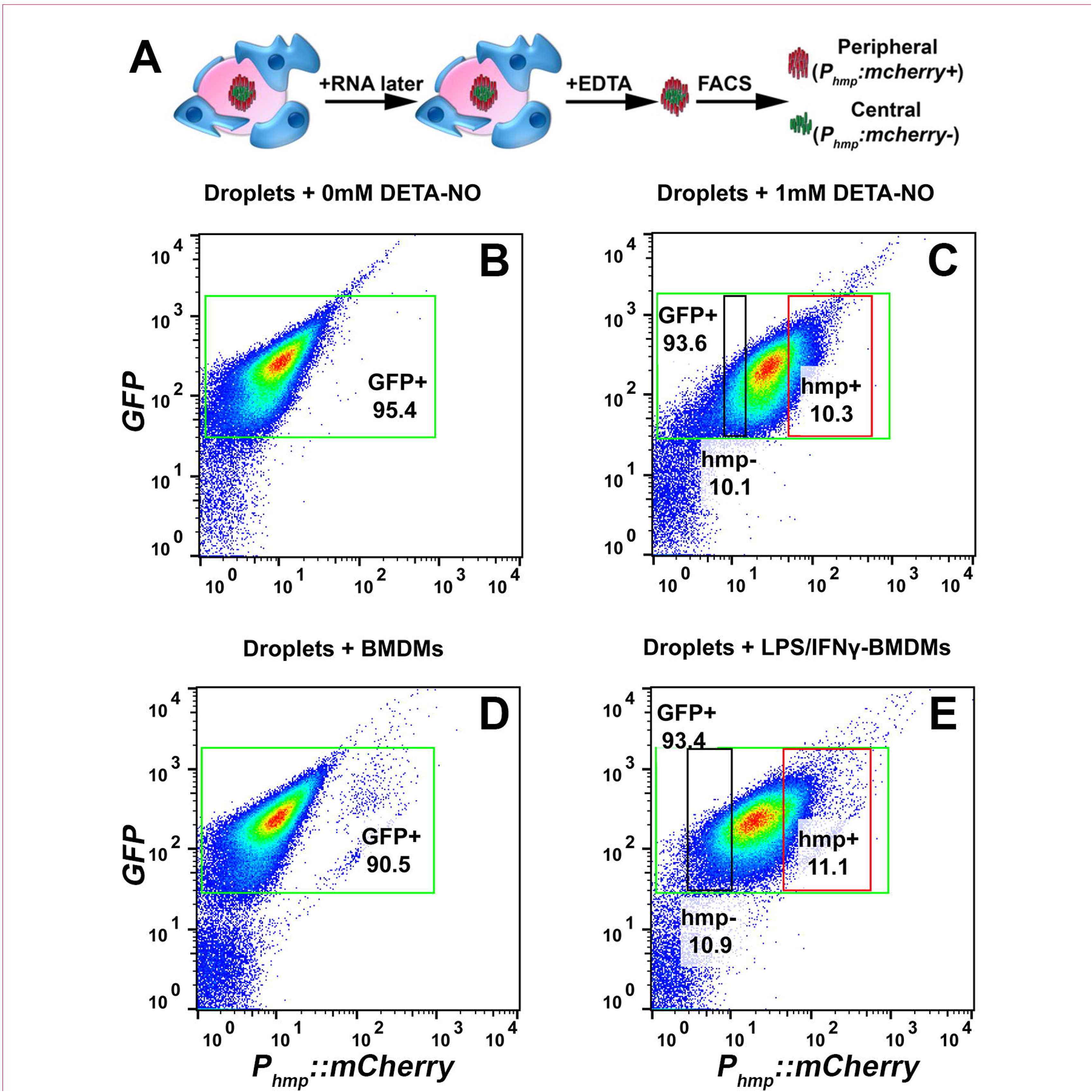
Flow sorting of *Yersinia pseudotuberculosis* subpopulations originating from alginate droplets. (A). Experimental overview. WT *Yptb gfp+, P_hmp::mcherry_* was encapsulated into alginate droplets and incubated for 7 hrs in DMEM to allow formation of microcolonies. The droplets were then incubated with either DETA-NONOate or BMDMs in DMEM for 4 hrs, and RNA*later* was added to the droplet-encompassed microcolonies. After subsequent extraction of *Y. pseudotuberculosis*, the bacterial subpopulations were sorted by flow cytometry into peripheral *hmp*^+^ mCherry(HI) and central *hmp*^-^ mCherry (LO) populations. (B-E) Flow cytometry of *Yptb* from droplets incubated with: (B) 0 mM DETA-NONOate; (C) 1 mM DETA-NONOate; (D) BMDMs; or (E) LPS/IFNγ-primed BMDMs. (B-E) Green rectangular gates: GFP+ events. (C,E) Red rectangular gates: mCherry-HI (*hmp*^+^) events. Black rectangular gates: mCherry-LO (*hmp*^-^) events. Flow was performed using 488nm (GFP) and 561 nm (mCherry) laser excitation, with GFP emission detected at 510-540 nm and mCherry detected at 602-627 nm emission windows.

### *Yptb* in the high *Phmp::mCherry* gate are transcriptionally distinct from other *Yptb*

Using the strategy described in Fig. 4, bacterial subpopulations were isolated, and differential RNAseq analysis comparing the *Phmp::mcherry*-HI gate to the *Phmp::mcherry*-LO gates was completed for 3 paired biological replicates of droplets that were exposed to either NONOate or IFN-activated BMDMs (Fig. 5) (38). Identical analysis was also performed for droplet-grown bacteria in the presence or absence of nonactivated-BMDMs. Among the most highly upregulated genes in response to macrophage droplet adherence were those associated with a subset of amino acid biosynthesis pathways, especially tryptophan and arginine (Figs. 5A,B). A similar starvation set was observed for peripherally localized bacteria exposed to IFNγ/LPS-activated BMDMs, although in the latter case, arginine starvation was no longer apparent (Fig. 5C,D; Supplemental Dataset 1). In contrast, amino acid biosynthesis genes were not upregulated in the presence of NONOate, arguing for macrophage specificity (Fig. 5E).

**Figure 5.**
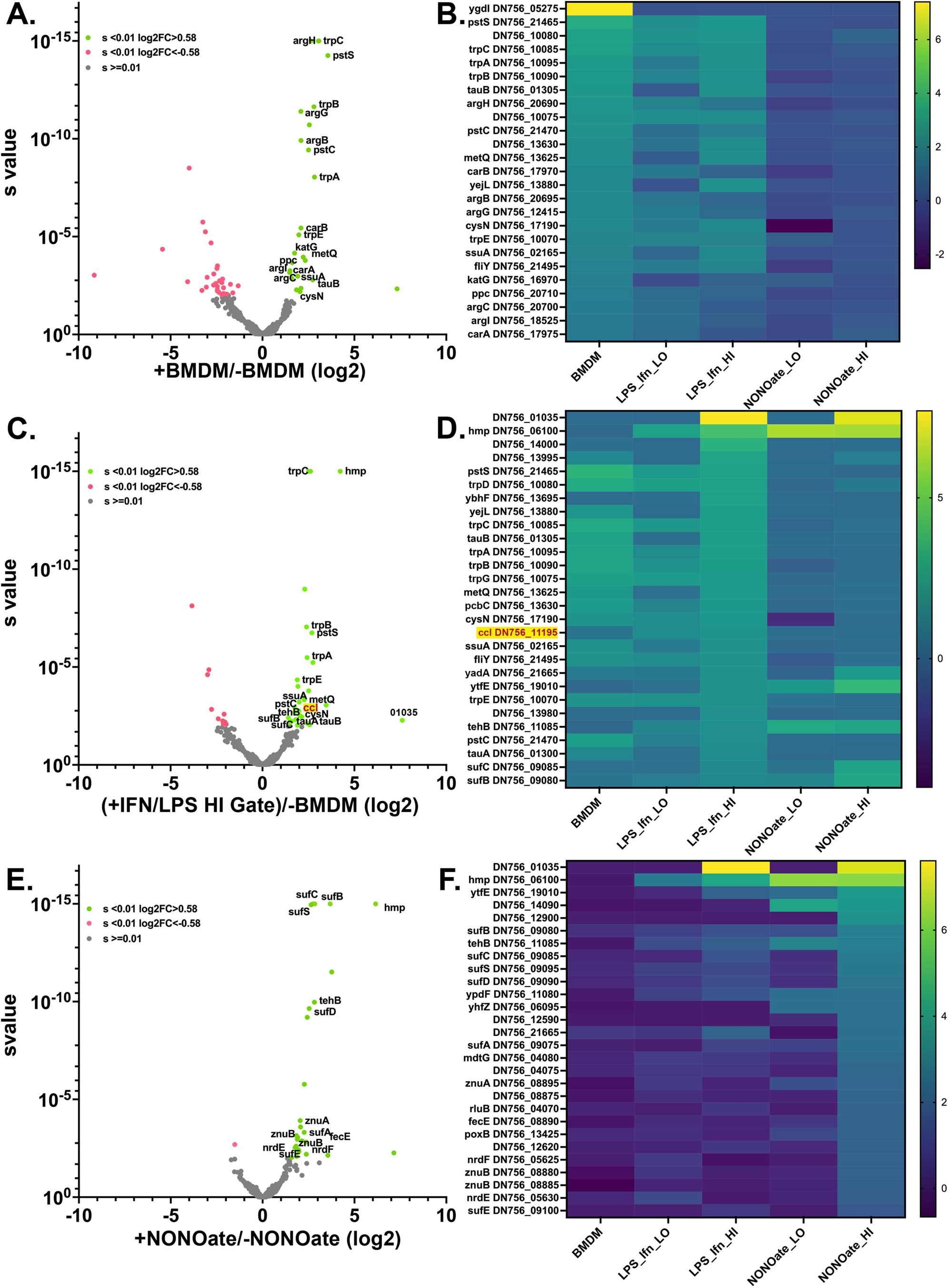
*Yptb* originating from the high *Phmp::mCherry* gate are transcriptionally distinct from the rest of the bacterial population. *Yptb* was grown in droplets in DMEM (Materials and Methods), exposed to noted conditions, RNA was isolated from sorted gates (Fig. 4), and RNA-tagseq analysis was performed using the noted conditions. (A,C,E) Volcano plots show statistical significance plotted as a function of the magnitude of differential expression for droplet-grown bacteria, comparing untreated droplets (-BMDM or -NONOate) to the noted bacteria populations. Red: s-value < 0.01 and log_2_-fold change of <-0.58; Gray: s-value ≥0.01; Green: s-value < 0.01 and log_2_-fold change of > 0.58. (B,D,F) Heat maps of log_2_ fold change relative to untreated droplets (-BMDM or -NONOate) are presented for 25-30 of the most highly upregulated genes when exposed to (B) unstimulated BMDMs, unsorted; (D) LPS/IFNγ-primed BMDMs, sorted *P_hmp_-mcherry*-HI; and (F) 1 mM NONOate, sorted for *P_hmp_-mcherry*-HI. Data are presented for the following populations: BMDM: exposed to unstimulated macrophages; LPS_Ifn_LO: exposed to LPS/IFNγ-primed BMDMs and sorted for *P_hmp_*-mCherry-LO fluorescence population; LPS_Ifn_HI: exposed to LPS/IFNγ-primed BMDMs and sorted for the *P_hmp_*-mCherry-HI fluorescence population; NONOate_LO: incubated with 1 mM NONOate and sorted for the *P_hmp_*-mCherry-LO low fluorescence population; NONOate_HI: incubated with 1 mM NONOate and sorted for the *P_hmp_*-mCherry-HI fluorescence population. Sorted populations defined in Fig. 4.

Bacterial microcolonies incubated with IFN/LPS-activated BMDMs showed a profile that was a hybrid of the responses to macrophages and to NONOate (Fig. 5C-F). Peripheral bacteria, gated for high mCherry expression after exposure to either activated BMDMs or NONOate displayed high expression levels of both *hmp* and the cysteine-rich protein-encoding DN756_01035, which is presumably a component of the NO detoxification response (Figs. 5D, F). In addition, genes encoding Fe-S cluster biosynthesis proteins SufA/B were upregulated relative to droplet-grown untreated controls in both peripherally localized populations (Fig. 5D, 5F, NONOate-HI). Interestingly, one of the few genes showing similar downregulation in response to activated BMDMs and NONOate was *frdA,* which encodes the Fe-S protein fumarate reductase, potentially reducing the size of Fe-S cluster center pools that need to be rejuvenated after NO attack (Supplemental Dataset 1). Most notably, there was a class of transcripts only induced in the presence of IFNγ/LPS-activated BMDMs and not by the other treatments. These included prophage-associated genes and a predicted ABC regulated efflux pump (*ybhF*, DN756_13695). Of the IFNγ/LPS-BMDM activated genes, induction of *ccl* (DN756_11195) was observed, which encodes a protein involved in degrading itaconate, a metabolite produced by macrophages in response to IFN treatment (Fig. 5D) (39).

When the response of the peripherally-localized bacteria to activated BMDM was compared to untreated macrophages, the unique set of genes upregulated in response to macrophage products comes into better focus (Fig. 6A; Supplemental Dataset 2). These included genes associated with protection from RNS (*tehB* (40)), and Fe-S cluster repair (*ytfE*) (41, 42), further indicating that NO is the primary driver of the response to activated macrophages. The lack of arginine starvation in the peripherally-localized bacteria noted above is supported by the lower expression of arginine biosynthesis-encoding genes (*carAB*, *argB*) relative to nonactivated controls (Fig. 6A), which is somewhat surprising, as macrophages can scavenge arginine to be used as the precursor of NO (43, 44). It is possible that a general reduction in metabolic rate in response to NO exposure suppresses transcription of arginine biosynthetic genes in peripherally localized bacteria. As a consequence, the specificity of the stress response to NO was notable in the peripherally localized bacteria.

**Figure 6.**
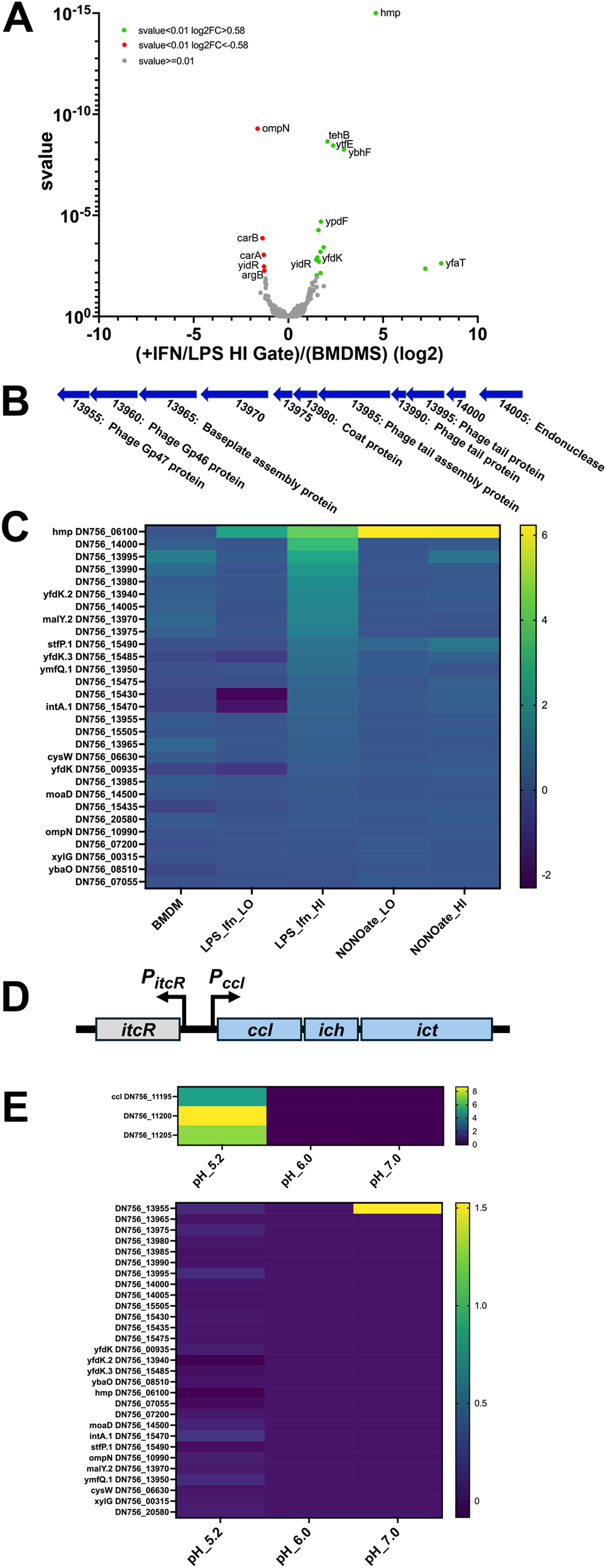
Spatial regulation of the response to activated BMDM. (A) Transcriptional response of *Yptb* located in the *P_hmp_-mcherry* high fluorescence gate (HI) compared to that after incubation of droplets with unstimulated BMDMs in DMEM. The statistical significance plotted as a function of the magnitude of differential expression of the bacteria isolated from the two gates. Red: s-value < 0.01 and log_2_-fold change of <-0.58; Gray: s-value ≥0.01; Green: s-value < 0.01 and log_2_-fold change of > 0.58. (B) Genetic map of cryptic phage locus harboring genes differentially upregulated in the *P_hmp_-mcherry* high fluorescence gate (HI) compared to untreated droplet-grown bacteria. (C). Heat map of the 29 transcripts, chosen based on their being the most highly differentially-transcribed genes comparing *P_hmp_-mcherry* high fluorescence gate (HI) to *P_hmp_-mcherry* low fluorescence gate (LO). BMDM: transcript expression after incubation with unstimulated macrophages compared to no macrophage incubation; LPS_Ifn_LO: exposed to LPS/IFNγ-primed BMDMs and sorted for *P_hmp_-mcherry*-LO population, compared to no macrophage incubation; LPS_Ifn_HI: exposed to LPS/IFNγ-primed BMDMs and sorted for the *P_hmp_-mcherry*-HI population compared to no macrophage incubation; NONOate_LO: incubated with 1 mM NONOate and sorted for the *P_hmp_-mcherry*-LO population compared to no NONOate incubation; NONOate_HI: incubated with 1 mM NONOate and sorted for the *P_hmp_-mcherry*-HI population compared to no NONOate incubation. Scale: log_2_ differences displayed. Sorted populations defined in Fig. 4. (D). Genetic map of itaconate degradation operon and ItcR regulator. (E). Phage genes are not induced by itaconate. *Yptb* was grown in broth and exposed to 1.0 mM itaconate as described at pH 5.2, 6.0, and 7.0 with log_2_ fold change shown compared to absence of NONOate at noted pH. Genes chosen were from panel B. Top Panel: *ccl* operon, the other most differentially regulated genes in the bottom panel.

Regarding genes not clearly associated with the NO response, the peripheral population showed downregulation of *ompN*, consistent with alterations in membrane integrity or permeability. In addition, there was induction of a series of genes predicted to encode phage structural proteins, perhaps due to induction of a *Yptb* prophage (Fig. 6B,C; compare HI gate to BMDM). Although prophages are known to be induced by DNA damage (45), there was no evidence for an SOS response preferentially in the HI gate (Supplemental Dataset 3). Furthermore, treatment with NONOate alone failed to result in induction of most of these genes (Fig. 6B,C; Supplemental Dataset 3). Therefore, a combination of macrophage exposure and NO production is associated with a noncanonical stress response that can activate prophage expression in these microcolonies. Together, these results indicate that NO is the predominant antibacterial activity produced by activated BMDMs located distant from microcolonies.

### The macrophage secreted metabolite itaconate is sensed by microcolonies

The induction of *ccl* was the most notable unexpected regulatory effect of exposure to activated macrophages (Fig. 5C,D). This gene encodes the first step in the pathway for degrading itaconate, a dicarboxylic acid produced by the interferon-regulated immunoresponsive gene 1 (*Irg1*) (39, 46), which has antimicrobial effects on several intracellular pathogens, especially in acidic conditions (47–50). *Yersinia* species are among a subset of bacteria that can degrade itaconate to bypass toxicity (51) by encoding a three-gene operon (Fig. 6D), which breaks down itaconate to pyruvate and acetyl-CoA, and is regulated by ItcR (51). Degradation of itaconate is necessary for *Y. pestis* survival in activated macrophages, but a role for restricting extracellular *Yersinia* within mammalian tissues has not been established (52).

As itaconate may drive differential expression of *Yptb* genes in the presence of IFN/LPS-activated BMDMs, we determined if upregulation of *Yptb* genes in the peripheral HI-mCherry population (Fig. 6C) could be a consequence of itaconate sensing (Fig. 6E). *Yptb* was cultured in triplicate broth-grown cultures in the presence or absence of itaconate using 3 different pH conditions, and RNAseq analysis was performed on the samples. Each of the three genes in the itaconate degradation operon (*ccl, icl, itc*) were upregulated in response to itaconate, but only at pH=5.2 conditions (Fig. 6E), as observed previously for itaconate signaling in multiple organisms (48, 51, 53). Except for DN756_13955, encoding a putative phage baseplate protein (Fig. 6B), there was no evidence for itaconate regulation of the genes upregulated in the presence of activated macrophages (Fig. 6E). Therefore, itaconate regulation appeared limited to the *ccl* cluster, while induction of prophage-related genes appeared to be a response to other stresses caused by IFN activated macrophages located in the vicinity of the microcolony.

Plasmid-based transcriptional reporters for itaconate degradation were constructed to analyze the dynamics of the itaconate response. These contained the intact *itcR* regulatory gene and a gene encoding a fluorescent protein controlled by the *ccl* promoter (Fig. 7A; Table 1) (54, 55). WT *Yptb* (Fig. 7) or a Δ*ccl* derivative (Supplemental Fig. S2) harboring the *P_ccl_*::*gfp* plasmid were grown in acidic conditions at pH = 6.0 or 5.2 (Fig. 7B, C), and fluorescence intensity was continuously monitored at 26°C. The addition of itaconate had little effect on the growth of *Yptb* at any pH, with post-exponential phase yields matching or exceeding those observed in the absence of itaconate (Fig. 7C). Consistent with itaconate control of similar operons (48, 51, 53), activation of the *ccl* promoter (*P_ccl_*) required low pH, with concentrations of 1.0 mM in broth adjusted to pH=5.2 providing near maximal induction when assayed in late exponential phase (Fig. 7D; 6 hr timepoint). When challenged with itaconate at pH=6.0, induction of the *P_ccl_*::*gfp* reporter required a high itaconate concentration (5.0 mM) and yielded lower maximal induction levels than observed at pH=5.2. Notably, peak induction occurred as the bacteria approached post-exponential phase, but there was little accumulation afterward, in contrast to the results at pH=5.2, where accumulation continued during post-exponential phase (Fig. 7C).

**Figure 7.**
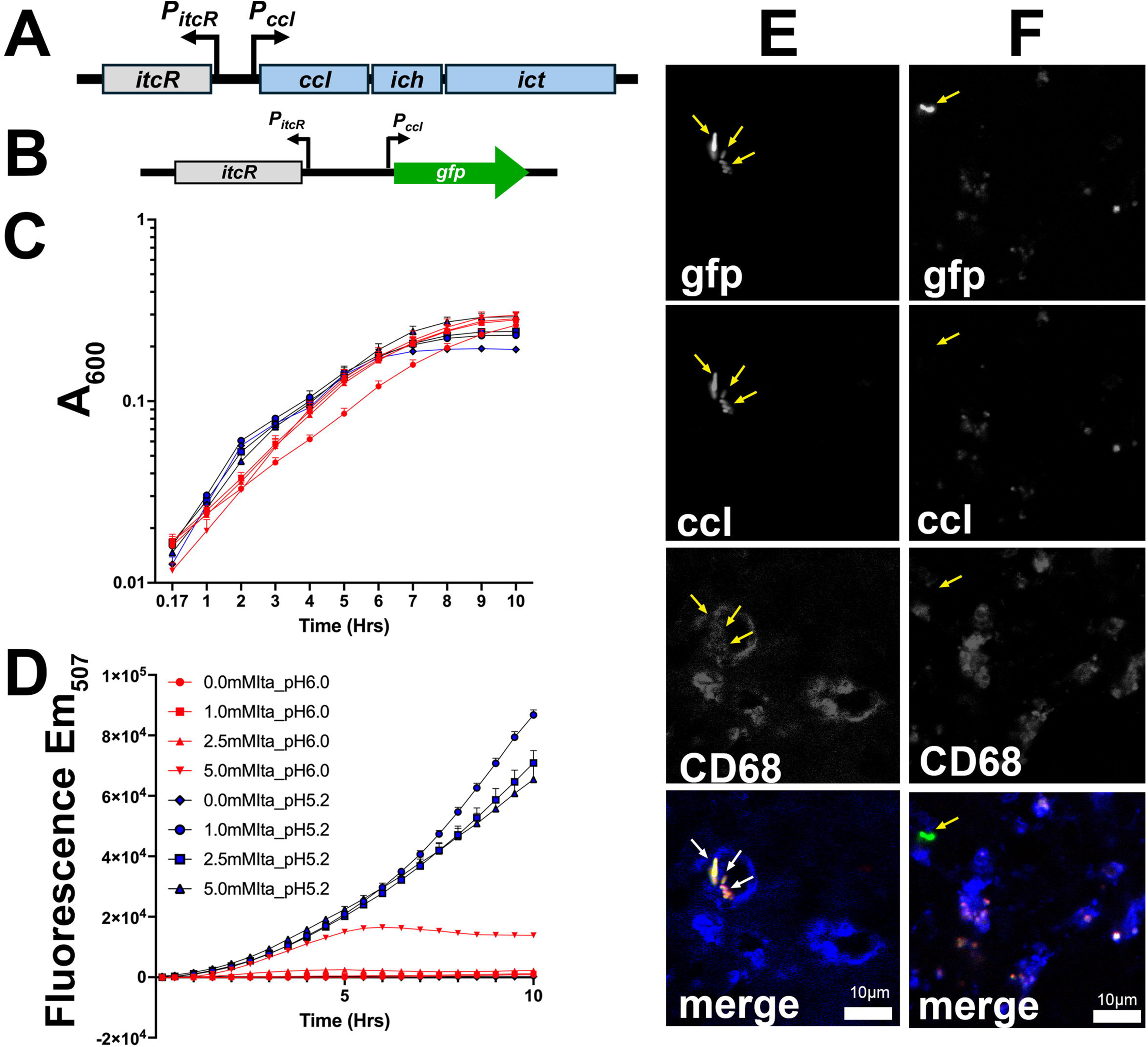
*Yptb* rarely experiences high itaconate concentrations in the spleen. (A,B). Fluorescent reporter plasmids that allow sensing of itaconate. Plasmids have intact *itcR* regulator that senses itaconate and controls *ccl* promoter. Fusions to *mcherry* and *gfp* were constructed as identical plasmids (Materials and Methods). (C,D). Overnight broth cultures of *Y. pseudotuberculosis* containing the *P_ccl_*_-_*gfp* reporter were diluted to A_600_=0.03 in LB adjusted to either pH=5.2 or pH=6.0 in the presence of noted itaconate concentrations. Increased mass was read at A_600_ (C), while fluorescence of the GFP reporter was measured excitation = 485/9 nm and emission = 507/9 nm (D). (E,F) C57BL/6 mice were injected intravenously with 10^3^ WT *gfp+* harboring the *itcR-p_ccl_:mcherry* reporter and spleens were harvested 3 days post inoculation. Spleen tissue was probed with α-CD68 (blue) and visualized by fluorescence microscopy. Arrows: bacterial cells. Scale bars: 10 μm.

**Table 1.**
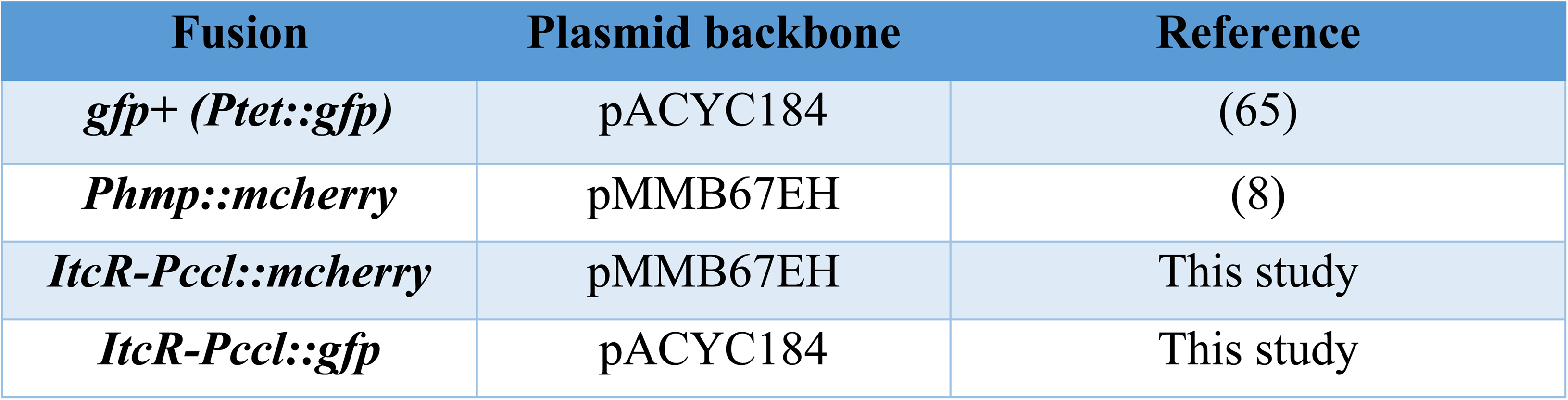
Plasmids used in this study.

As acidic conditions were necessary to observe induction of the *P_ccl_* reporter, we investigated if itaconate penetrates a *Yptb* microcolony during infection using the fluorescent reporter system. To this end, mice were injected intravenously with 10^3^ *Yptb gfp+* harboring *ItcR-Pccl:mcherry* itaconate reporter. Three days post-infection, spleens were removed and processed for fluorescence microscopy. Only a few microcolonies showed induction at this time point (< 10% of the colonies scanned). Itaconate has been primarily studied as a product of activated macrophages, although there are reports of neutrophil production of the metabolite(56), so we determined if induction could result from rare close encounter with macrophages by probing tissues with α-CD68 (macrophage marker) followed by fluorescence microscopy. Macrophage-associated *Yptb* showed expression of the itaconate reporter (Fig. 7E), arguing that itaconate can penetrate a microcolony only after rare direct contact with macrophages in the tissue microenvironment.

### The itaconate degradation operon and microcolony formation during *Yptb* growth in deep tissue sites

We predicted that neutrophils surrounding microcolonies may serve as a buffer layer preventing high levels of macrophage-secreted itaconate from penetrating the microcolony. To determine if the itaconate degradation operon contributes to survival of *Yptb* during infection, and if neutrophils buffer from macrophage attack, mice were inoculated intravenously with *Yptb* WT(pGFP^+^) or *Δccl-ich-ict* (pGFP^+^) in the presence or absence of neutrophil depletion. Neutrophil depletion should result in macrophages directly contacting each microcolony and increasing itaconate exposure. Three days post-infection, spleens were harvested to quantify CFU and visualize microcolonies using fluorescence microscopy. Challenge with the *Yptb*Δ(*ccl-ich-ict*) strain resulted in a small decrease in CFU relative to WT in mock-depleted mice (Fig. 8A). Neutrophil depletion had little effect on this result and, in fact, any differences between WT and the Δ(*ccl-ich-ict*) strain did not rise to the level of significance (Fig. 8A,C; Mann-Whitney test). Microcolony size was then determined by image analysis of sections, using GFP fluorescence (Fig. 8E). There was a small shift downward in microcolony size of the (*ccl-ich-ict*) strain after neutrophil depletion, which was not observed in the mock-depleted animals (Fig. 8B,D). Although specific for the absence of neutrophils, the shift was not significant based on multiple statistical tests (unpaired t test; Mann-Whitney).

**Figure 8.**
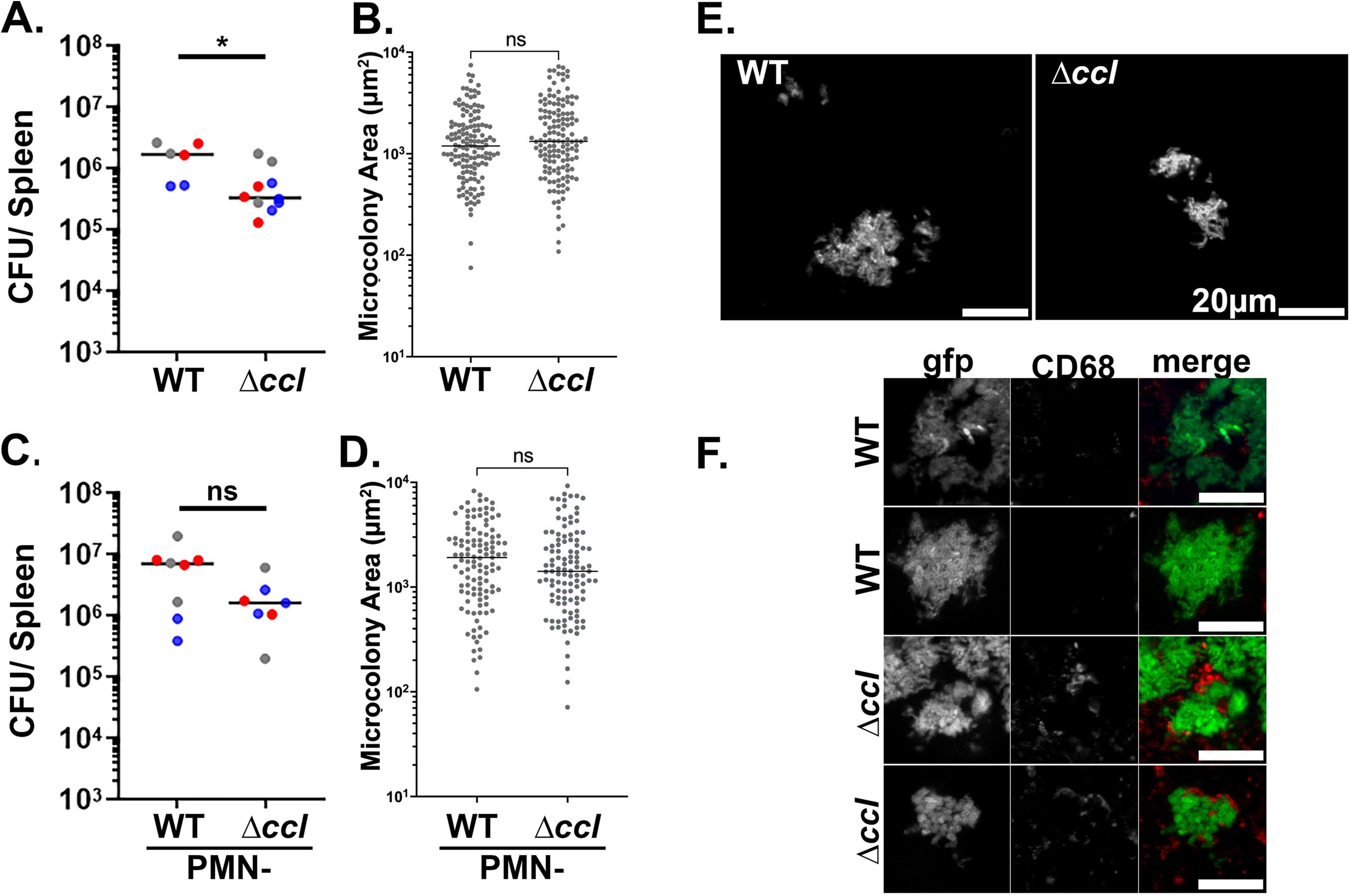
Effects of loss of itaconate degradation operon during tissue growth of *Yptb*. C57BL/6 mice were intraperitoneally injected with α-Ly6G (clone 1A8) or PBS 16 hours prior to and 24 hours post-infection to deplete or perform mock-depletion of neutrophils (PMN). Mice were inoculated intravenously (i.v.) with 10^3^ WT *gfp+* or *Δccl-ich-ict gfp+* bacteria, and spleens were harvested 3 days post-inoculation (PI) for quantification of CFU and fluorescence microscopy. Lines represent the median. (A, B): Mock-depleted C57BL/6 mice. (A) Total CFU and (B) microcolony areas were quantified (Materials and Methods). Lines represent the median. (C,D): α-Ly6G depleted C57BL/6 mice. (C) Total CFU and (D) microcolony areas were quantified. Lines represent the median. Blue, red, and gray dots: Individual mice from 3 separate experiments. Statistics: Mann-Whitney test, **p<0.005, *p<0.05, ns: not significant. (E,F):Representative WT and *Δccl-ich-ict* microcolonies showing variation in morphology. Scale bar: 20 μm (E): Representative WT and *Δccl-ich-ict* microcolonies after infection in neutrophil-depleted mice. Scale bar: 20 μm

We noted that microcolonies formed by *Yptb Δccl-ich-ict* appeared to have aberrant morphology and, in general, lacked rod shape, indicating a stress response (Fig. 8F; examples in Supplemental Fig. 3). Interestingly, the bloated bacteria appeared to be in contact with macrophages (Fig. 8F). We then analyzed images of both WT and *Δccl-ich-ict* after challenge of either neutrophil-replete or depleted animals. Evidence of bloated bacteria was not correlated with either the presence of neutrophils or the loss of the itaconate degradation pathway (Supplemental Fig. 3). Rather, colonies containing 20 or fewer bacteria were often found to harbor bloated organisms, consistent with disrupted colony formation being associated with aberrant bacterial morphology. Samples enriched for smaller colonies, therefore, are more likely to accumulate bloated bacterial cells. These results point to the model that extracellular growth largely tolerates the absence of the itaconate degradation pathway even when in contact with macrophages in tissues.

## Discussion

Invasive bacterial pathogens spread to distant tissue sites and establish areas where they can multiply to high numbers despite the presence of host innate immune cells. Growth within tissues is followed by the recruitment of neutrophils, macrophages, and inflammatory monocytes. These inflammatory sites can lead to granulomas or abscesses when immune cells fail to clear the infection (4, 5). There are few culture models that accurately replicate the complex structure of pathogen and host immune cells growing in tissue. To address this gap, we previously developed a 3D model to reconstruct *Yptb* inflammatory sites (18).

During *Yptb* spread into the murine spleen, neutrophils are unable to limit the bacterial burden, leading to recruitment of macrophages and other inflammatory monocytes to the outside of the neutrophil layer (18). Our microfluidics-based droplet strategy closely models this second step in the process, in which macrophages are called to respond to the absence of bacterial clearance. We view the gel microdroplet as topologically mimicking the zone between the microcolony and the macrophages, which is inhabited by neutrophils during the formation of a *Yptb* microcolony in tissues (8). In support of this hypothesis, both the NO generator DETA-NONOate and activated BMDMs drove a bacterial transcriptional response that was morphologically indistinguishable from that observed in tissues (Fig. 3) (8). The ability of macrophages in their activated state to modulate bacterial gene expression is likely due to the production of soluble antimicrobial mediators because there is no direct contact with the *Yptb* microcolony. A variety of small secretory products of macrophages have been identified, but their roles in controlling bacterial growth or modulating microbial gene expression are greatly understudied (57). Consistent with that interpretation that macrophage metabolites control bacterial gene expression by action-at-a-distance in this system, we see little evidence for cell contact-activated gene expression. In our previous work (3, 8), we demonstrated that Yop expression is induced only at the extreme periphery of the microcolony, where bacteria contact neutrophils. The lack of such contact, and the presence of Ca^2+^ in the medium, suppresses expression of Yops above what is observed at 37°C during growth in culture.

To complete RNA-seq on *Yptb* subpopulations, the previously developed agarose-containing gel system had to be reworked and replaced with alginate as the droplet matrix. Both the microcolony morphology and the spatial regulation of the *Phmp* reporter in response to either DETA-NONOate or activated BMDMs were similar to that observed in agarose droplets. The use of this strategy provided immediate benefits and enabled estimation of the amount of RNI generated by activated macrophages surrounding the microcolonies. Based on the flow cytometry data of the *hmp* reporter under the two conditions, the amount of RNI produced by activated BMDMs, which is presumably similar to that observed in tissues, is higher than that produced by 1 mM DETA-NONOate,.

We hypothesized that *Yptb* growing in the presence of macrophages generates peripheral and central subpopulations that have distinct transcriptional profiles. RNA-seq analysis revealed that the bacterial subpopulation with high *Phmp* reporter firing showed a response that was largely a hybrid of that observed after droplet contact with macrophages and that observed after exposure to DETA-NONOate. In addition to Hmp induction, genes associated with iron-cluster formation as well as a poorly annotated cysteine-rich protein that could act as a sink for toxic RNI-byproducts were observed in the presence of both DETA-NONOate and activated macrophage adherence. This indicates that NO is the dominant factor produced by activated BMDMs that target *Yptb* from a distance. Hmp detoxifies NO into nitrate, so we might have expected that Hmp would generate a considerable pool of nitrate. There was no evidence that *Yptb* utilizes nitrate as an alternative electron acceptor despite the availability of this source, with no induction of nitrate reductase (Fig. 5; Supplemental Dataset 1). This is consistent with oxygen not being limited and remaining the preferred electron acceptor. There was similar lack of evidence for alternate electron acceptors during microcolony growth in the murine spleen (8).

There were two notable additions to this hybrid response, involving the induction of the itaconate degradation pathway (*ccl*) and prophage genes. Although not specific for the peripheral subpopulation (Fig. 5D), *ccl* upregulation did require macrophage activation by LPS/IFNγ, presumably in response to liberation of itaconate, which is synthesized within macrophages by the well-characterized interferon-upregulated Irg1 gene. The encoded enzyme catalyzes the conversion of the TCA cycle product aconitate to the dicarboxylate itaconate (39). Originally thought to solely operate as a down modulator of the macrophage inflammatory response (39), the discovery that the *Y. pestis ccl* cluster encoding itaconate degradation is required for intracellular survival within activated macrophages indicates that the metabolite plays a role in restricting intracellular bacterial growth of this pathogen that is closely related to *Yptb* (51, 52). Work in *L. pneumophila* (58, 59) and *S. typhimurium* (53) supports the concept for restriction of intracellular bacteria, particularly in acidic environments. There is also evidence that itaconate liberated by activated macrophages is sensed by extracellular pathogens, such as *Pseudomonas aeruginosa*, resulting in membrane stress to the microbe (60). Furthermore, degradation of itaconate is associated with successful persistent infection by *Pseudomonas* (51, 60, 61).

Intravenous infections with a *Yptb* strain deleted for the *ccl-ich-ict* operon argued that itaconate degradation plays, at best, a subtle role in supporting *Yptb* proliferation in deep tissue sites. There were small reductions in bacterial yields and microcolony size within spleen tissues, and there was only rare firing of a *ccl-mcherry* reporter in tissue microcolonies. Those rare microcolonies did appear to directly abut macrophages, indicating that transcription of the itaconate degradation operon was linked to close contact with NO-producing cells (Fig. 8). Given the role of acidic growth conditions both in activating the operon and restricting growth in the presence of itaconate, it seems unlikely that itaconate could contribute to effective growth restriction extracellularly. Rather, the most likely route for growth restriction by itaconate would be targeting of rare, internalized bacteria within an acidic compartment in activated macrophages, similar to that observed for *Y. pestis* (52). Induction of the operon by extracellular itaconate could provide *Yptb* with a preparatory induction signal that allows protection once the bacteria are internalized. As the *Yptb* type III secretion system primarily acts to prevent phagocytic internalization of the bacteria, this strategy presumably acts as a back-up system to protect bacteria that fail to block internalization.

The second notable addition to the macrophage/NO hybrid response was the activation of prophage-associated transcripts after exposure to activated macrophage (Fig. 6). There was clear spatial regulation to these transcripts, as they were found primarily associated with the peripherally localized bacterial population. Prophage induction is usually associated with stress responses to DNA damage or reactive oxygen species (ROS) (62). There was little evidence of a transcriptional response to ROS or DNA damage, or signatures of an SOS response (45), while the exposure of bacteria to DETA-NONOate or itaconate alone was insufficient to reproduce the prophage transcript activation (Fig. 6D). We think it possible the combination of byproducts generated by macrophage metabolism and RNI may cause some low-level stress response that results in prophage induction and mimics something akin to a DNA-damage response.

Overall, the alginate droplet system is a powerful technique that allows us to analyze bacterial subpopulations within a microcolony and identify additional macrophage-secreted mediators that target *Yptb* (Fig. 9). RNA-seq revealed that there is a *Yptb* subpopulation within microcolony undergoing a specific stress response resulting in prophage induction (Fig. 9). Furthermore, the itaconate degradation operon*, ccl-ich-ict,* was upregulated throughout the colony exposed to activated BMDMs, indicating that itaconate can penetrate these bacteria (Fig. 9). Future work will involve incorporating neutrophils to allow the construction of a complete inflammatory site that recapitulates tissue behavior. This emphasizes the advantages of using alginate as a matrix, because the polymer is easily tunable via alginate lyase, allowing control over the size of the matrix surrounding the microcolony. Theoretically, neutrophils can be brought into contact with microcolonies by controlled degradation (63) and, furthermore, this microfluidic approach could serve as a blueprint for dissecting spatial transcriptomics and complex host-pathogen interactions in a number of other bacterial pathogens

**Figure 9.**
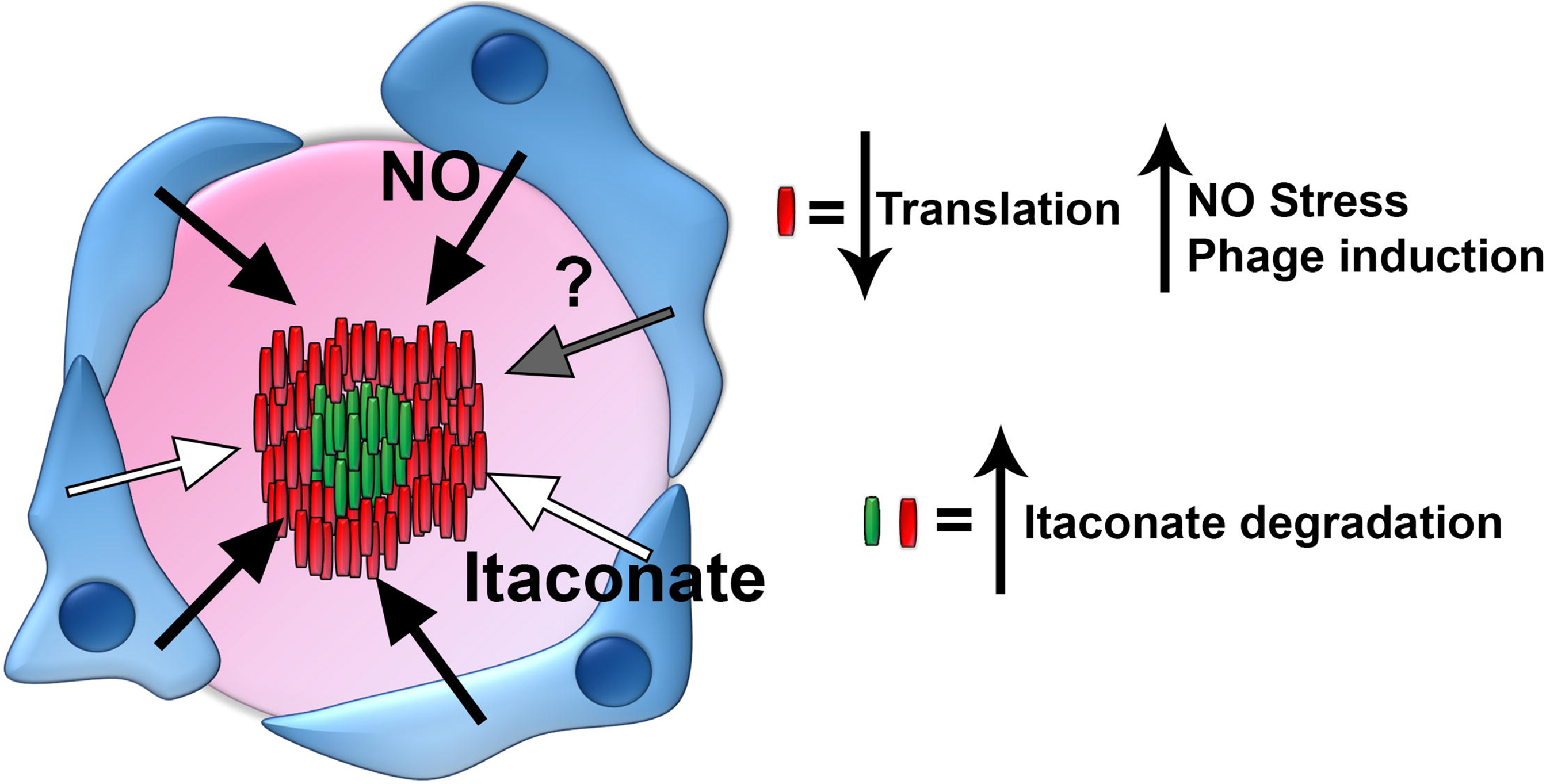
Bioengineered droplets allow for the transcriptional analysis of bacterial subpopulations and inform about the tissue microenvironment. Bacteria are grown *in situ* within alginate droplets prior to challenge with iNOS^+^ BMDMs. Activated BMDMs secrete NO, itaconate, and other metabolites which target a *Yptb* microcolony. Hmp+ bacteria on the periphery of a microcolony upregulate genes involved in iron sulfur cluster assembly and phage induction. This indicates that the peripheral cells within a microcolony are likely slow-growing. Throughout the entire colony, itaconate is sensed.

## Materials and Methods

### Bacterial strains & growth conditions

Plasmids used are described in Table 1. All experiments used derivatives of the *Y. pseudotuberculosis* strain IP2666 (Table 2; (64)). For all droplet experiments, bacteria were grown overnight into stationary phase in 2xYT broth ((65); (8) at 26 °C with rotation. For itaconate (Sigma # I29204-100G) exposure in broth cultures, overnight cultures were diluted 1:100 in LB and rotated at 37 °C for 4 hours in the presence of noted concentrations of itaconate.

**Table 2.**
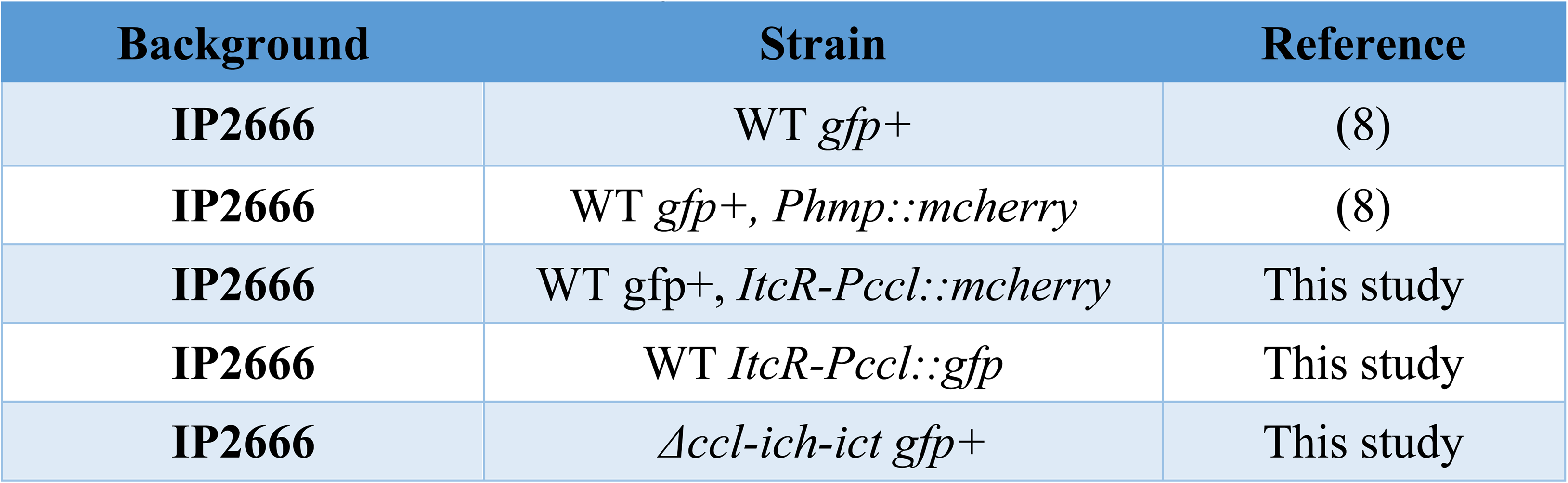
Strains used in this study.

### Generation of plasmid-based reporter strains

The *Y. pseudotuberculosis P_hmp_::mCherry* reporter strain in this study has been previously described (8, 65). GFP^+^ strains encode GFP under the control of the unrepressed *P_tet_* promoter of pACYC184 (66). The *P_hmp_::mCherry* transcriptional fusion was constructed by fusing the *hmp* promoter to *mCherry* using overlap extension PCR, and cloning into the pMMB67EH plasmid (67), Strains that harbor both reporters are co-transformants of the compatible pACYC184 and pMMB67EH plasmids. The *ItcR-P_ccl_::mcherry* and *ItcR-P_ccl_::gfp* transcriptional fusions were constructed by fusing *ItcR-ccl* promoter to *mCherry* and *gfp* using Gibson Assembly^®^ (New England BioLabs) and cloning into pMMB67EH and pACYC184, respectively (54, 55).

### Generation of ccl-ich-ict mutant strains

Strains having deletions in the gene of interest were generated by amplifying the start codon + 3 downstream codons, the 3’ terminal 3 codons + the stop codon and fusing these fragments to generate a 24 base pair scar sequence. Deletion constructs were amplified with 500 base pairs flanking sequence on each side, cloned into the suicide vector, pSR47s (68, 69), transformed into DH5α λ*pir* and mated into *Y*.

*pseudotuberculosis* IP2666 using the helper strain RK600 (65). Sucrose counterselection was used to isolate strains that had recombined the *ccl-ich-ict* construct and removed the WT copy of the gene, then strains were confirmed by PCR and sequencing of PCR product.

### Droplet generation

CAD-designed microfluidics chips having 38 independent devices controlled by two ports were constructed according to published protocols (70) at the Boston College Nanofabrication Facility (https://www.bc.edu/bc-web/schools/mcas/sites/cleanroom.html). Prior to droplet generation, the device was primed with fluorinated oil (3M Novec 7500) and connected to two programmable syringe pumps (Harvard Apparatus 11 Elite Series) located in a temperature-controlled room at 37 °C. One pump was fitted with a 1ml syringe (BD #309628) having a 27-gauge needle (BD #305109), which was filled with 1.5% Pico-Surf 1 (Dolomite Microfluidics, 5% in Novec 7500) in Novec 7500 oil (18). In parallel, overnight cultures were diluted 1:2000 in 1% Na-alginate (Fisher #AC177772500) in DMEM with glutamine, 10% FBS, 25mM Hepes and 25 mM CaCO_3_ nanoparticles (SSNano #1951RH) containing RGD-alginate at a concentration that was 5% of unmodified Na-alginate (Novamatrix, Novatach MVG GRGDSP), yielding approximately 5 × 10^6^ cells/ml prior to droplet generation. The mix was loaded into a 1 ml syringe that was fitted onto a second pump. Polyethylene (PE/2) tubing (SCI COM #BB31695-PE/2) was attached to each syringe and then inserted into the appropriate ports on the microfluidics device (18). The pump holding the oil-phase was set to 500 μl/hour while the other pump was set to 200 μl/hour and droplets were collected in a 1.5 ml microfuge tube through tubing that had been inserted into the droplet collection port.

### Oil Removal

100 μl droplets was introduced into a 1.5 ml microfuge tube and 400 μl of 1.5% Pico-Surf 1 in Novec 7500 oil was added to the droplets. To crosslink the alginate, 500 μl of 0.1% acetic acid in Novec 7500 oil was added to the droplets. After mixing by inverting the tube several times, the dilute acetic acid in oil was removed from the tube and 500 μl of 10% 1H,1H,2H,2H-Perfluoro-1-octanol (PFO) (Sigma #370533-25G) in Novec 7500 was added to the droplets. After mixing vigorously for approximately 10 seconds, 500 μl of DMEM with 10% FBS was added and pipetted up and down to resuspend the droplets without disturbing the oil layer. The DMEM/droplet layer was transferred to a new 1.5 ml microfuge tube, subjected to centrifugation for 30 seconds at 250 RCF, and washed with DMEM. The remaining droplets were washed and resuspended in 2xYT broth supplemented with 5 mM CaCl_2_ to maintain alginate droplet integrity.

### Droplet dispersal

Droplets were washed once in DMEM at the indicated timepoints, and 900 μl of RNA*later* (Invitrogen #AM7021) was added to the samples dispersing droplets. Samples were incubated overnight at 4°C, then resuspended with 10 mL ice cold PBS. Samples were centrifuged at 5,000 x *g* for 15 minutes. The supernatant was removed and the bacterial pellets were resuspended in 500 μl of PBS sample, then prepared for flow analysis.

### Analysis of the nitric oxide response

To analyze the response of colonies to exogenous NO, droplets containing *Y. pseudotuberculosis* were generated, oil was removed, and the droplets were rotated for 7 hours at 26 °C in 2xYT broth to allow colony formation. After the 7-hour growth period, droplets were subjected to centrifugation for 30 seconds at 250 RCF and washed twice in DMEM. 50 μl of droplets were resuspended in 1 ml DMEM with glutamine (Gibco) supplemented with 10% FBS in 12-well non-tissue culture treated plates and exposed to DETA-NONOate (Sigma #AC32865) during growth at 37 °C with 5% CO_2_ for the indicated times.

### Analysis of the itaconate response

For each condition, 3 independent cultures were made from single colonies and grown overnight in LB broth with aeration at 26°C. The bacteria were then diluted 1:100 in LB broth and grown in same medium to A_600_ = 0.3. The bacteria were pelleted, washed three times in 1X PBS/Mg/Ca, and diluted 1:30 into DMEM/25 mM HEPES + 10% FBS equilibrated to noted pH levels in the presence or absence of 1 mM itaconate and prewarmed to 37°C. Bacteria were grown to A_600_ =0.1 and 50% of the cultures were pelleted and flash-frozen in liquid N_2_. The rest of the cultures were grown to A_600_ =0.3, pelleted and flash-frozen in liquid N_2_. RNA was extracted and processed for RNA-tagseq analysis.

### Isolation of bone marrow-derived macrophages

Bone marrow cells were isolated from femurs, tibias, and humeri of female 6-8-week-old C57BL/6 mice, and terminally differentiated into macrophages in medium containing mouse CSF generated as previously described (18, 71). Bone marrow-derived macrophages (BMDMs) were subsequently frozen in fetal bovine serum (FBS) with 10% dimethyl sulfoxide (DMSO) and plated one day prior to incubation with droplets in 90% fresh medium (DMEM with glutamine (Gibco) supplemented with 10% FBS) spiked with 10% sterile medium containing macrophage colony stimulating factor (MCSF) in 5% CO_2_ at 37°C. Medium containing MCSF was conditioned by growth in the presence of mouse 3T3 fibroblasts that overproduce mouse MCSF at 37 °C with 5% CO2, as previously described (71) (72). To stimulate BMDMs to induce iNOS expression, cells from frozen stocks were plated on non-tissue culture treated plates and allowed to recover overnight. The following day, BMDMs were stimulated with 200 U/ml IFNγ (Peprotech) and 100 ng/ml LPS (Sigma) for 12-15 hours (18, 35).

### Droplet-BMDM experiments

Droplet-BMDM experiments were completed as previously described (18). Droplets containing *Y. pseudotuberculosis* were generated, oil was removed, and the droplet-encased bacteria were grown with aeration for 7 hours at 26 °C in 2xYT broth. After the 7-hour growth period, droplets were pelleted for 30 seconds at 250 RCF, washed twice in DMEM supplemented with 10% FBS and supernatant was removed. 50 μl of droplets were transferred to a well in a 12-well non-tissue culture treated plate, approximately 2 × 10^6^ BMDMs in DMEM with 10% FBS were added to the well, and the incubation proceeded at 37 °C in the presence of 5% CO_2_ for 4 hours. Samples were fixed in 2% paraformaldehyde (PFA) in PBS, washed in PBS, and fluorescent reporters were visualized by microscopic observation.

### Murine model of systemic infection

Six to 8-week old female C57BL/6 mice obtained from Jackson Laboratories (Bar Harbor, ME) were injected intravenously with 10^3^ bacteria. Three days post inoculation, spleens were removed and processed as described previously (8, 65). For neutrophil depletions, mice were intraperitoneally injected with 50 μg rat anti-mouse Ly6G antibody (BD Biosciences # 551459) 16 hours prior to and 24 hours post-infection (34). All animal studies were approved by the Institutional Animal Care and Use Committee of Tufts University.

### Flow cytometry and fluorescence-activated cell sorting (FACS)

After microcolony dispersal, samples were filtered through 40 μm filters. Filtered samples were added to 5ml FACS tubes containing 2 ml PBS on ice. Fluorescent reporters were detected using a Bio-Rad S3e cell sorter. Purity Sort Mode was used during sorting, and the priority was set to the high *mCherry* subpopulation when applicable. Samples were sorted into 5 ml tubes containing 0.5ml ice cold PBS. 3-10 ×10^6^ events were collected for each sample. After sorting, samples were subjected to centrifugation 5,000 x g for 15 min at 4°C, supernatants were removed, and pellets were overlayed with 400 uL of ice cold PBS. Samples were flash frozen in liquid nitrogen and stored at −80°C for later use.

### RNA extraction

200 μl of pre-heated (95 °C) Max Bacterial Enhancement reagent (Ambion #16122-012) was added to frozen bacterial pellets. Afterward, samples were transferred to a 1.5 ml microfuge tube and incubated at 95 °C for 5 minutes. Extraction of total RNA was conducted using the Zymo Direct-Zol Microprep Kit and following the protocol for cells in suspension (Zymo #R2026).

### RNA-seq sample preparation

cDNA libraries were generated following the RNAtag-seq protocol as previously described (73, 74), but ribosomal RNA (rRNA) was depleted using Illumina® Ribo-Zero Plus rRNA depletion kit to deplete prokaryotic rRNA and any contaminating eukaryotic rRNA. Libraries were sequenced at High-Output for 75 cycles on Illumina NextSeq500 (TUCF Core).

### Computational Analysis

RNA-Seq data was analyzed using customized pipeline. In brief, raw reads were demultiplexed by 5’ and 3’ indices and mapped to the reference genomes (CP032566.1 and CP032567.1) using bwa mem (75). Mapped reads were aggregated by featureCount (76), and differential expression was calculated with DESeq2 (38, 76). *apeglm* estimation [https://doi.org/10.1093/bioinformatics/bty895] was used within the lfcShrink function. In each pair-wise differential expression comparison, significant differential expression was filtered based on svalue < 0.01. Hierarchical cluster analysis was performed for the top 50 genes having the most significant differential expression.

### Fluorescence microscopy

After 4 hours of nitric oxide exposure or BMDM challenge, droplets were fixed in 2% PFA in PBS for 10 minutes. Droplets were visualized by transferring the sample to a microscope slide and covering with a coverslip. To analyze tissue samples, C57BL/6 mice were inoculated intravenously with the indicated strains. Three days post inoculation, spleens were harvested and fixed in 4% PFA in PBS for 3 hours, tissue was frozen-embedded in Sub Xero freezing media (Mercedes Medical) and cut into 10 μm sections using a cryostat microtome (Microm HM505E). To visualize reporters, sections were thawed in PBS, stained with Hoechst at a 1:10,000 dilution, washed in PBS, and coverslips were mounted using ProLong Gold (Life Technologies). For immunofluorescence staining of macrophages, sections were thawed, blocked with 2% BSA, permeabilized with PBT (0.2% Triton X-100, 0.1% BSA), and stained using a 1:1000 dilution of Alexa Fluor^®^ 594 anti-mouse CD68 antibody (Biolegend # 137020). Tissue was imaged with a 63x objective on a Zeiss Axio Observer.Z1 (Zeiss) microscope with Colibri.2 LED light source, an Apotome 2 (Zeiss) for optical sectioning, and an ORCA-R^2^ digital CCD camera (Hamamatsu) (8, 65).

### Determination of microcolony area

To determine microcolony area in tissue, images were captured by fluorescence microscopy using a 20x lens with analysis in Volocity^TM^ as previously described (18). Microcolony area was determined by identifying a threshold that defines edges of microcolonies in the GFP channel and determining the number of pixels in the region of interest (ROI). The data were then converted to metric scale using a stage micrometer and displayed as μm^2^.

### Quantitative image analysis

MATLAB scripts were written to calculate the average intensity of fluorescence about the periphery of microcolonies and point of lowest fluorescence as previously described (18); copy archived at https://github.com/elifesciences-publications/droplet.

### Deposition of RNA-seq data

All raw RNA-seq datasets have been deposited in SRA under BioProject PRJNA1120393.

## Acknowledgements

We would like to thank Dr. Albert Tai and members of the Tufts University Core Facility (TUCF) Genomics unit for instruction, help, and guidance throughout this work, as well as Mr. Steve Kwok of the Tufts Thorley-Lawson Sorting Facility for hands-on help with these studies. We thank Drs. Bixi He, Kevin Manera, Atish Roy Chowdhury, and Efrat Hamami for review of the manuscript. The work in this study was supported by NIAID grants R01AI110684 and R21 151593 to RRI and U01AI124302 and R01AI110724 to TvO. SC was supported by predoctoral training grant T32 TM007310 from NIGMS

## References

1. Carter PB, Collins FM. 1974. The route of enteric infection in normal mice. J Exp Med 139:1189–203.

2. Cheng AG, Kim HK, Burts ML, Krausz T, Schneewind O, Missiakas DM. 2009. Genetic requirements for *Staphylococcus aureus* abscess formation and persistence in host tissues. FASEB J 23:3393–404.

3. Simonet M, Richard S, Berche P. 1990. Electron microscopic evidence for in vivo extracellular localization of *Yersinia pseudotuberculosis* harboring the pYV plasmid. Infect Immun 58:841–5.

4. Cheng AG, DeDent AC, Schneewind O, Missiakas D. 2011. A play in four acts: *Staphylococcus aureus* abscess formation. Trends Microbiol 19:225–32.

5. Pagan AJ, Ramakrishnan L. 2018. The Formation and Function of Granulomas. Annu Rev Immunol 36:639–665.

6. Zhang Y, Khairallah C, Sheridan BS, van der Velden AWM, Bliska JB. 2018. CCR2(+) Inflammatory Monocytes Are Recruited to *Yersinia pseudotuberculosis* Pyogranulomas and Dictate Adaptive Responses at the Expense of Innate Immunity during Oral Infection. Infect Immun 86.

7. Peterson LW, Philip NH, DeLaney A, Wynosky-Dolfi MA, Asklof K, Gray F, Choa R, Bjanes E, Buza EL, Hu B, Dillon CP, Green DR, Berger SB, Gough PJ, Bertin J, Brodsky IE. 2017. RIPK1-dependent apoptosis bypasses pathogen blockade of innate signaling to promote immune defense. J Exp Med 214:3171–3182.

8. Davis KM, Mohammadi S, Isberg RR. 2015. Community behavior and spatial regulation within a bacterial microcolony in deep tissue sites serves to protect against host attack. Cell Host Microbe 17:21–31.

9. Barnes PD, Bergman MA, Mecsas J, Isberg RR. 2006. *Yersinia pseudotuberculosis* disseminates directly from a replicating bacterial pool in the intestine. J Exp Med 203:1591–601.

10. Peterson ST, Dailey KG, Hullahalli K, Sorobetea D, Matsuda R, Sewell J, Yost W, O’Neill R, Bobba S, Apenes N, Sherman ME, Balazs GI, Assenmacher CA, Cox A, Lanza M, Shin S, Waldor MK, Brodsky IE. 2025. TNF signaling maintains local restriction of bacterial founder populations in intestinal and systemic sites during oral *Yersinia* infection. mBio 16:e0177925.

11. Black DS, Bliska JB. 2000. The RhoGAP activity of the *Yersinia pseudotuberculosis* cytotoxin YopE is required for antiphagocytic function and virulence. Mol Microbiol 37:515–27.

12. Durand EA, Maldonado-Arocho FJ, Castillo C, Walsh RL, Mecsas J. 2010. The presence of professional phagocytes dictates the number of host cells targeted for Yop translocation during infection. Cell Microbiol 12:1064–82.

13. Rosqvist R, Forsberg A, Wolf-Watz H. 1991. Intracellular targeting of the *Yersinia* YopE cytotoxin in mammalian cells induces actin microfilament disruption. Infect Immun 59:4562–9.

14. Viboud GI, Bliska JB. 2005. Yersinia outer proteins: role in modulation of host cell signaling responses and pathogenesis. Annu Rev Microbiol 59:69–89.

15. Mejia E, Bliska JB, Viboud GI. 2008. *Yersinia* controls type III effector delivery into host cells by modulating Rho activity. PLoS Pathog 4:e3.

16. Schotte P, Denecker G, Van Den Broeke A, Vandenabeele P, Cornelis GR, Beyaert R. 2004. Targeting Rac1 by the *Yersinia* effector protein YopE inhibits caspase-1-mediated maturation and release of interleukin-1beta. J Biol Chem 279:25134–42.

17. Songsungthong W, Higgins MC, Rolan HG, Murphy JL, Mecsas J. 2010. ROS-inhibitory activity of YopE is required for full virulence of *Yersinia* in mice. Cell Microbiol 12:988–1001.

18. Clark SA, Thibault D, Shull LM, Davis KM, Aunins E, van Opijnen T, Isberg R. 2020. Topologically correct synthetic reconstruction of pathogen social behavior found during *Yersinia* growth in deep tissue sites. Elife 9.

19. Andersen T, Auk-Emblem P, Dornish M. 2015. 3D Cell Culture in Alginate Hydrogels. Microarrays (Basel) 4:133–61.

20. Chen Q, Utech S, Chen D, Prodanovic R, Lin JM, Weitz DA. 2016. Controlled assembly of heterotypic cells in a core-shell scaffold: organ in a droplet. Lab Chip 16:1346–9.

21. Hunt NC, Hallam D, Karimi A, Mellough CB, Chen J, Steel DHW, Lako M. 2017. 3D culture of human pluripotent stem cells in RGD-alginate hydrogel improves retinal tissue development. Acta Biomater 49:329–343.

22. Tan WH, Takeuchi S. 2007. Monodisperse Alginate Hydrogel Microbeads for Cell Encapsulation. Advanced Materials 19:2696–2701.

23. Bruchet M, Melman A. 2015. Fabrication of patterned calcium cross-linked alginate hydrogel films and coatings through reductive cation exchange. Carbohydr Polym 131:57–64.

24. Velasco D, Tumarkin E, Kumacheva E. 2012. Microfluidic encapsulation of cells in polymer microgels. Small 8:1633–42.

25. Utech S, Prodanovic R, Mao AS, Ostafe R, Mooney DJ, Weitz DA. 2015. Microfluidic Generation of Monodisperse, Structurally Homogeneous Alginate Microgels for Cell Encapsulation and 3D Cell Culture. Adv Healthc Mater 4:1628–33.

26. Cai S, Zhao M, Fang Y, Nishinari K, Phillips GO, Jiang F. 2014. Microencapsulation of *Lactobacillus acidophilus* CGMCC1.2686 via emulsification/internal gelation of alginate using Ca-EDTA and CaCO3 as calcium sources. Food Hydrocolloids 39:295–300.

27. Zhao M, Qu F, Cai S, Fang Y, Nishinari K, Phillips GO, Jiang F. 2015. Microencapsulation of Lactobacillus acidophilus CGMCC1.2686: Correlation Between Bacteria Survivability and Physical Properties of Microcapsules. Food Biophysics 10:292–299.

28. Cha BH, Shin SR, Leijten J, Li YC, Singh S, Liu JC, Annabi N, Abdi R, Dokmeci MR, Vrana NE, Ghaemmaghami AM, Khademhosseini A. 2017. Integrin-Mediated Interactions Control Macrophage Polarization in 3D Hydrogels. Adv Healthc Mater 6.

29. Giancotti FG, Ruoslahti E. 1999. Integrin signaling. Science 285:1028–32.

30. Hoesli CA, Kiang RLJ, Raghuram K, Pedroza RG, Markwick KE, Colantuoni AMR, Piret JM. 2017. Mammalian Cell Encapsulation in Alginate Beads Using a Simple Stirred Vessel. J Vis Exp doi:10.3791/55280.

31. Hu Y, Mao AS, Desai RM, Wang H, Weitz DA, Mooney DJ. 2017. Controlled self-assembly of alginate microgels by rapidly binding molecule pairs. Lab Chip 17:2481–2490.

32. Khavari A, Nyden M, Weitz DA, Ehrlicher AJ. 2016. Composite alginate gels for tunable cellular microenvironment mechanics. Sci Rep 6:30854.

33. Frawley ER, Karlinsey JE, Singhal A, Libby SJ, Doulias P-T, Ischiropoulos H, Fang FC. 2018. Nitric Oxide Disrupts Zinc Homeostasis in *Salmonella enterica* Serovar Typhimurium. mBio 9:e01040–18.

34. Green ER, Clark S, Crimmins GT, Mack M, Kumamoto CA, Mecsas J. 2016. Fis Is Essential for *Yersinia pseudotuberculosis* Virulence and Protects against Reactive Oxygen Species Produced by Phagocytic Cells during Infection. PLoS Pathog 12:e1005898.

35. Mosser DM, Gonçalves R. 2015. Activation of Murine Macrophages, p 14.2.1-14.2.10, Current Protocols in Immunology doi:10.1002/0471142735.im1402s111.

36. Nuss AM, Beckstette M, Pimenova M, Schmuhl C, Opitz W, Pisano F, Heroven AK, Dersch P. 2017. Tissue dual RNA-seq allows fast discovery of infection-specific functions and riboregulators shaping host-pathogen transcriptomes. Proc Natl Acad Sci U S A 114:E791–E800.

37. Yang C, Frei H, Rossi FM, Burt HM. 2009. The differential in vitro and in vivo responses of bone marrow stromal cells on novel porous gelatin-alginate scaffolds. J Tissue Eng Regen Med 3:601–14.

38. Love MI, Huber W, Anders S. 2014. Moderated estimation of fold change and dispersion for RNA-seq data with DESeq2. Genome Biol 15:550.

39. Michelucci A, Cordes T, Ghelfi J, Pailot A, Reiling N, Goldmann O, Binz T, Wegner A, Tallam A, Rausell A, Buttini M, Linster CL, Medina E, Balling R, Hiller K. 2013. Immune-responsive gene 1 protein links metabolism to immunity by catalyzing itaconic acid production. Proc Natl Acad Sci U S A 110:7820–5.

40. Sebbane F, Lemaitre N, Sturdevant DE, Rebeil R, Virtaneva K, Porcella SF, Hinnebusch BJ. 2006. Adaptive response of *Yersinia pestis* to extracellular effectors of innate immunity during bubonic plague. Proc Natl Acad Sci U S A 103:11766–71.

41. Bodenmiller DM, Spiro S. 2006. The *yjeB* (*nsrR*) gene of *Escherichia coli* encodes a nitric oxide-sensitive transcriptional regulator. J Bacteriol 188:874–81.

42. Justino MC, Almeida CC, Goncalves VL, Teixeira M, Saraiva LM. 2006. *Escherichia coli* YtfE is a di-iron protein with an important function in assembly of iron-sulphur clusters. FEMS Microbiol Lett 257:278–84.

43. Sakuma I, Stuehr DJ, Gross SS, Nathan C, Levi R. 1988. Identification of arginine as a precursor of endothelium-derived relaxing factor. Proc Natl Acad Sci U S A 85:8664–7.

44. Marletta MA, Yoon PS, Iyengar R, Leaf CD, Wishnok JS. 1988. Macrophage oxidation of L-arginine to nitrite and nitrate: nitric oxide is an intermediate. Biochemistry 27:8706–11.

45. Witkin EM. 1975. Elevated mutability of polA derivatives of *Escherichia coli* B/r at sublethal doses of ultraviolet light: evidence for an inducible error-prone repair system (“SOS repair”) and its anomalous expression in these strains. Genetics 79 Suppl:199–213.

46. Cordes T, Wallace M, Michelucci A, Divakaruni AS, Sapcariu SC, Sousa C, Koseki H, Cabrales P, Murphy AN, Hiller K, Metallo CM. 2016. Immunoresponsive Gene 1 and Itaconate Inhibit Succinate Dehydrogenase to Modulate Intracellular Succinate Levels. J Biol Chem 291:14274–14284.

47. Cordes T, Michelucci A, Hiller K. 2015. Itaconic Acid: The Surprising Role of an Industrial Compound as a Mammalian Antimicrobial Metabolite. Annu Rev Nutr 35:451–73.

48. Wang H, Fedorov AA, Fedorov EV, Hunt DM, Rodgers A, Douglas HL, Garza-Garcia A, Bonanno JB, Almo SC, de Carvalho LPS. 2019. An essential bifunctional enzyme in *Mycobacterium tuberculosis* for itaconate dissimilation and leucine catabolism. Proc Natl Acad Sci U S A 116:15907–15913.

49. Nguyen TV, Alfaro AC, Young T, Green S, Zarate E, Merien F. 2019. Itaconic acid inhibits growth of a pathogenic marine *Vibrio* strain: A metabolomics approach. Sci Rep 9:5937.

50. Duncan D, Lupien A, Behr MA, Auclair K. 2021. Effect of pH on the antimicrobial activity of the macrophage metabolite itaconate. Microbiology (Reading) 167.

51. Sasikaran J, Ziemski M, Zadora PK, Fleig A, Berg IA. 2014. Bacterial itaconate degradation promotes pathogenicity. Nat Chem Biol 10:371–7.

52. Pujol C, Grabenstein JP, Perry RD, Bliska JB. 2005. Replication of *Yersinia pestis* in interferon gamma-activated macrophages requires *ripA*, a gene encoded in the pigmentation locus. Proc Natl Acad Sci U S A 102:12909–14.

53. Chen M, Sun H, Boot M, Shao L, Chang SJ, Wang W, Lam TT, Lara-Tejero M, Rego EH, Galan JE. 2020. Itaconate is an effector of a Rab GTPase cell-autonomous host defense pathway against *Salmonella*. Science 369:450–455.

54. Hanko EKR, Minton NP, Malys N. 2018. A Transcription Factor-Based Biosensor for Detection of Itaconic Acid. ACS Synth Biol 7:1436–1446.

55. Hanko EKR, Minton NP, Malys N. 2019. Design, cloning and characterization of transcription factor-based inducible gene expression systems. Methods Enzymol 621:153–169.

56. Tomlinson KL, Riquelme SA, Baskota SU, Drikic M, Monk IR, Stinear TP, Lewis IA, Prince AS. 2023. *Staphylococcus aureus* stimulates neutrophil itaconate production that suppresses the oxidative burst. Cell Rep 42:112064.

57. Nathan CF. 1987. Secretory products of macrophages. J Clin Invest 79:319–26.

58. Naujoks J, Tabeling C, Dill BD, Hoffmann C, Brown AS, Kunze M, Kempa S, Peter A, Mollenkopf HJ, Dorhoi A, Kershaw O, Gruber AD, Sander LE, Witzenrath M, Herold S, Nerlich A, Hocke AC, van Driel I, Suttorp N, Bedoui S, Hilbi H, Trost M, Opitz B. 2016. IFNs Modify the Proteome of *Legionella-*Containing Vacuoles and Restrict Infection Via IRG1-Derived Itaconic Acid. PLoS Pathog 12:e1005408.

59. Price JV, Russo D, Ji DX, Chavez RA, DiPeso L, Lee AY, Coers J, Vance RE. 2019. IRG1 and Inducible Nitric Oxide Synthase Act Redundantly with Other Interferon-Gamma-Induced Factors To Restrict Intracellular Replication of *Legionella pneumophila*. mBio 10.

60. Riquelme SA, Liimatta K, Wong Fok Lung T, Fields B, Ahn D, Chen D, Lozano C, Saenz Y, Uhlemann AC, Kahl BC, Britto CJ, DiMango E, Prince A. 2020. Pseudomonas aeruginosa Utilizes Host-Derived Itaconate to Redirect Its Metabolism to Promote Biofilm Formation. Cell Metab 31:1091–1106 e6.

61. Runtsch MC, O’Neill LAJ. 2020. *Pseudomonas* Persists by Feeding off Itaconate. Cell Metab 31:1045–1047.

62. DeMarini DM, Lawrence BK. 1992. Prophage induction by DNA topoisomerase II poisons and reactive-oxygen species: role of DNA breaks. Mutat Res 267:1–17.

63. Sakai S, Inagaki H, Inamoto K, Taya M. 2012. Wrapping tissues with a pre-established cage-like layer composed of living cells. Biomaterials 33:6721–7.

64. Balada-Llasat JM, Mecsas J. 2006. *Yersinia* has a tropism for B and T cell zones of lymph nodes that is independent of the type III secretion system. PLoS Pathog 2:e86.

65. Crimmins GT, Mohammadi S, Green ER, Bergman MA, Isberg RR, Mecsas J. 2012. Identification of MrtAB, an ABC transporter specifically required for *Yersinia pseudotuberculosis* to colonize the mesenteric lymph nodes. PLoS Pathog 8:e1002828.

66. Chang AC, Cohen SN. 1978. Construction and characterization of amplifiable multicopy DNA cloning vehicles derived from the P15A cryptic miniplasmid. Journal of Bacteriology 134:1141–1156.

67. Fiirstea J, Pansegrau W, Frank R, Blbcker H, Scholz P, Bagdasarian M, Lanka E. 1986. Molecular cloning of the plasmid RP4 primase region in a multi-host-range tacP expression vector. Gene 48:119–131.

68. Miller VL, Mekalanos JJ. 1988. A novel suicide vector and its use in construction of insertion mutations: osmoregulation of outer membrane proteins and virulence determinants in *Vibrio cholerae* requires *toxR*. J Bacteriol 170:2575–83.

69. Andrews HL, Vogel JP, Isberg RR. 1998. Identification of linked *Legionella pneumophila* genes essential for intracellular growth and evasion of the endocytic pathway. Infect Immun 66:950–8.

70. Mazutis L, Gilbert J, Ung WL, Weitz DA, Griffiths AD, Heyman JA. 2013. Single-cell analysis and sorting using droplet-based microfluidics. Nat Protoc 8:870–91.

71. Auerbuch V, Golenbock DT, Isberg RR. 2009. Innate immune recognition of *Yersinia pseudotuberculosis* type III secretion. PLoS Pathog 5:e1000686.

72. Leber JH, Crimmins GT, Raghavan S, Meyer-Morse NP, Cox JS, Portnoy DA. 2008. Distinct TLR- and NLR-mediated transcriptional responses to an intracellular pathogen. PLoS Pathog 4:e6.

73. Shishkin AA, Giannoukos G, Kucukural A, Ciulla D, Busby M, Surka C, Chen J, Bhattacharyya RP, Rudy RF, Patel MM, Novod N, Hung DT, Gnirke A, Garber M, Guttman M, Livny J. 2015. Simultaneous generation of many RNA-seq libraries in a single reaction. Nat Methods 12:323–5.

74. Jensen PA, Zhu Z, van Opijnen T. 2017. Antibiotics Disrupt Coordination between Transcriptional and Phenotypic Stress Responses in Pathogenic Bacteria. Cell Rep 20:1705–1716.

75. Li H, Durbin R. 2009. Fast and accurate short read alignment with Burrows-Wheeler transform. Bioinformatics 25:1754–60.

76. Liao Y, Smyth GK, Shi W. 2014. featureCounts: an efficient general purpose program for assigning sequence reads to genomic features. Bioinformatics 30:923–30.

