## Supplementary figures and images for "The transcriptional response of *Yersinia pseudotuberculosis* to macrophage-released metabolites during growth within synthetic microcolonies"

### Supplemental Figs

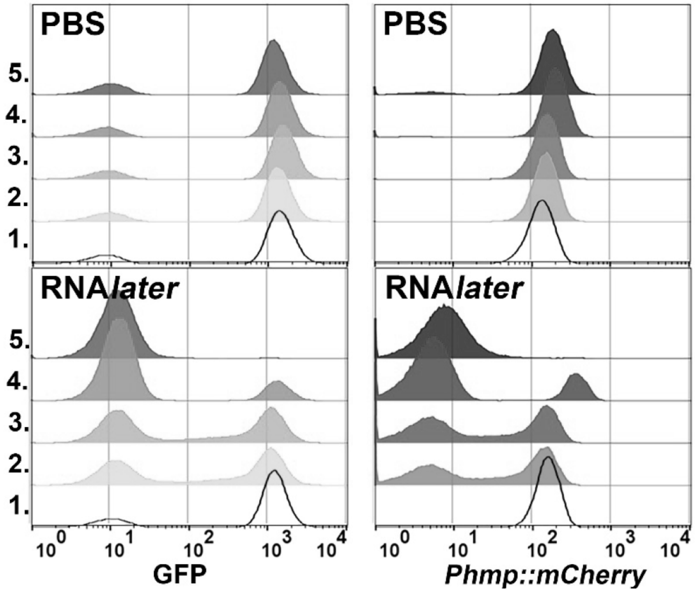

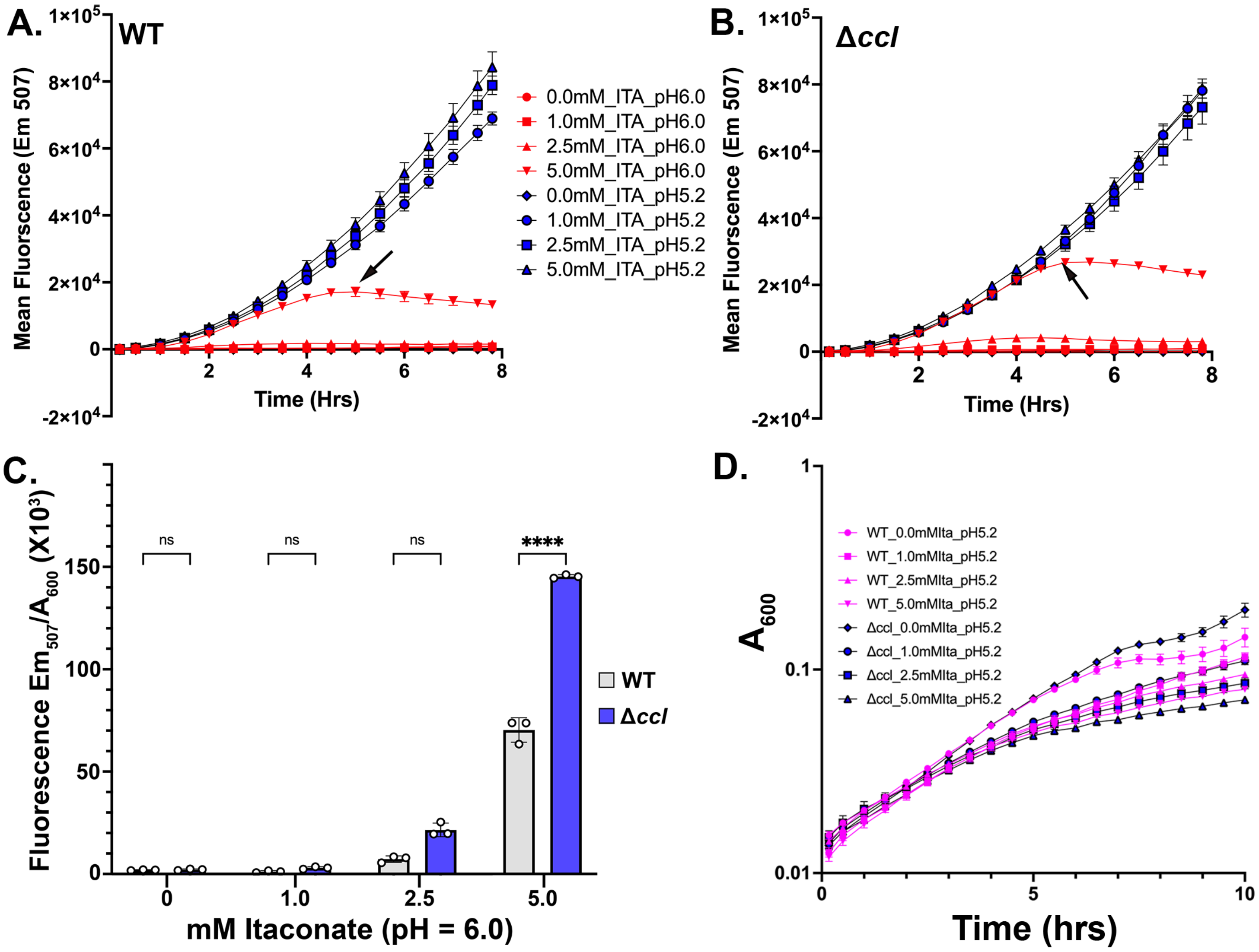

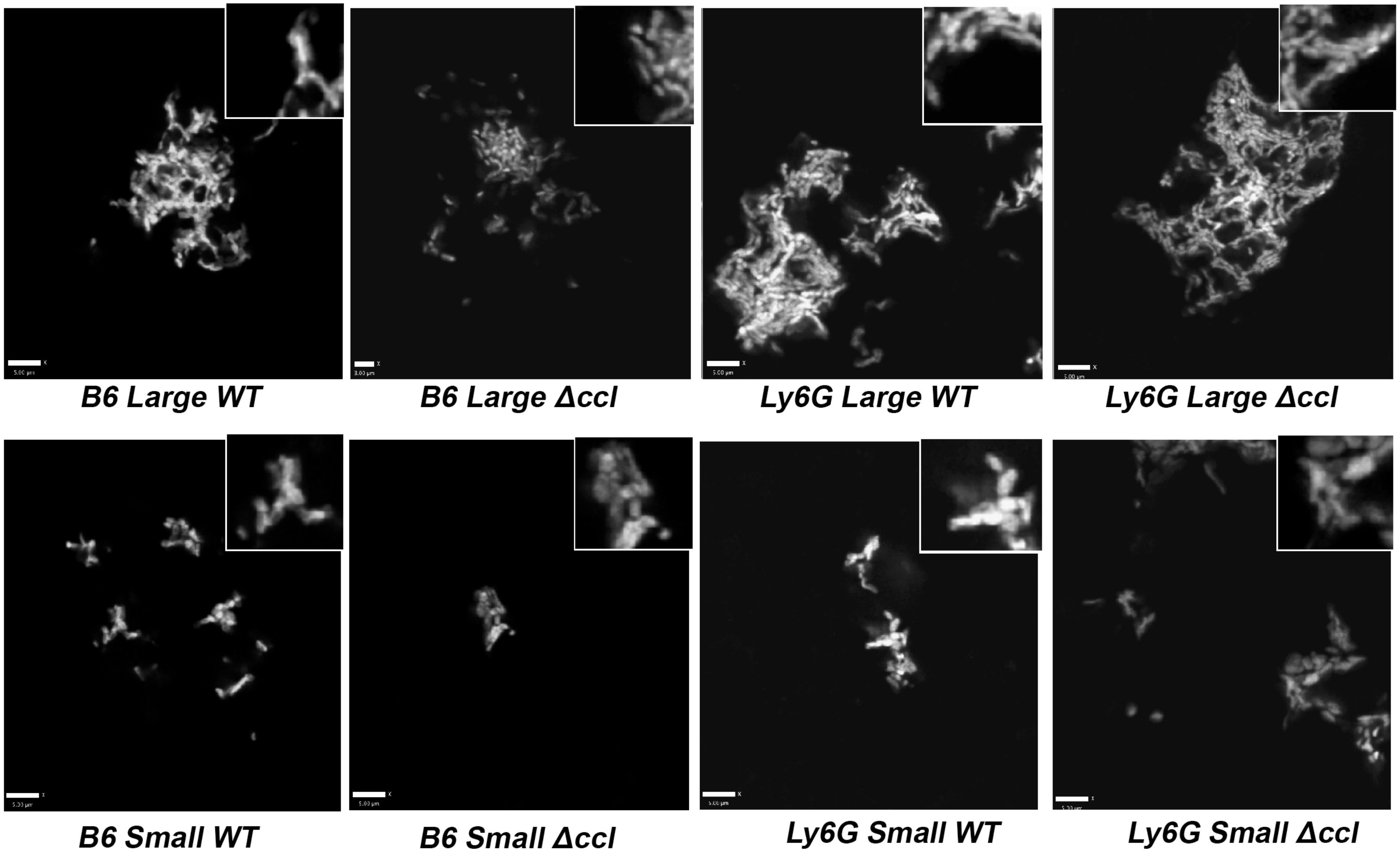
